# Correlative 3D microscopy of single cells using super-resolution and scanning ion-conductance microscopy

**DOI:** 10.1101/2020.11.09.374157

**Authors:** Vytautas Navikas, Samuel M. Leitao, Kristin S. Grussmayer, Adrien Descloux, Barney Drake, Klaus Yserentant, Philipp Werther, Dirk-Peter Herten, Richard Wombacher, Aleksandra Radenovic, Georg E. Fantner

## Abstract

High-resolution live-cell imaging is necessary to study complex biological phenomena. Modern fluorescence microscopy methods are increasingly combined with complementary, label-free techniques to put the fluorescence information into the cellular context. The most common high-resolution imaging approaches used in combination with fluorescence imaging are electron microscopy and atomic-force microscopy (AFM), originally developed for solid-state material characterization. AFM routinely resolves atomic steps, however on soft biological samples, the forces between the tip and the sample deform the fragile membrane, thereby distorting the otherwise high axial resolution of the technique. Here we present scanning ion-conductance microscopy (SICM) as an alternative approach for topographical imaging of soft biological samples, preserving high axial resolution on cells. SICM is complemented with live-cell compatible super-resolution optical fluctuation imaging (SOFI). To demonstrate the capabilities of our method we show correlative 3D cellular maps with SOFI implementation in both 2D and 3D with self-blinking dyes for two-color high-order SOFI imaging. Finally, we employ correlative SICM/SOFI microscopy for visualizing actin dynamics in live COS-7 cells with subdiffractional resolution.

## Introduction

Imaging living cells in vitro is crucial in deciphering the biochemical mechanisms underlying complex cellular activity such as cell motility^1^, differentiation^2^, membrane trafficking^3^, and cell-to-cell communication^4^. Our knowledge about the ultra-structure of a cell is almost exclusively derived from high-resolution electron microscopy (EM) on fixed and sectioned cells. Modern EM techniques can routinely reach sub-nm resolution and even provide the atomic structures of macromolecular complexes such as ribosomes^5^ or lately even whole viruses^6^. The function of the structures is typically inferred from the presence of specific molecules of interest by performing targeted expression or immunolabelling, followed by fluorescence microscopy techniques thus resulting in correlative light and electron microscopy (CLEM)^7^. Axial information can be obtained from EM images through serial sectioning^8^, however obtaining unperturbed 3D information about the shape of the cell membrane remains difficult and low throughput.

The portfolio of fluorescence imaging techniques has been expanded with multiple so-called super-resolution microscopy techniques, that circumvent the traditional diffraction limit of optical microscopy^9^. Many of these techniques trade-off temporal resolution for lateral resolution, and often require high light intensities causing light-induced cell damage in live cells^10^. An approach named super-resolution optical fluctuation imaging (SOFI)^11^ was developed to mitigate the negative phototoxic effects of long-term imaging by increasing acquisition speeds. SOFI relies on stochastic temporal fluctuations of the signal in independently blinking emitters and is less sensitive to varying fluorophore density and different blinking conditions, compared to localization microscopy approaches^12^. SOFI was demonstrated as a powerful imaging method which goes well beyond the diffraction limit and can be extended for live-cell 3D imaging^13^ or combined with self-blinking dyes^14^, simplifying the image acquisition pipeline.

On the other hand, scanning probe microscopy (SPM) can obtain nanometer resolution images of the 3D surface of unlabeled living cells in its physiological environment^15^. It has been proven as a versatile tool for biological imaging, and is often combined with fluorescence microscopy methods^16^. The most commonly used SPM technique, atomic-force microscopy (AFM), enables the study of the mechanobiology of fixed or living cells^17^, to image biological membranes^18^ and even track the dynamics of molecular assembly processes^19^. Label-free AFM methods combined with label-specific super-resolution fluorescence microscopy tools can provide maps of single-cells in unprecedented detail not only in fixed, but also in living samples^20,21^. AFM relies on the direct physical interaction with a sample, which usually deforms sensitive biological specimens and causes height artifacts in live-cell imaging^22^. To perform truly non-contact imaging in physiological conditions a scanning modality based on ionic current sensing was developed a few decades ago^23^ and was significantly improved with further developments for robust live-cell imaging^24^. Scanning-ion conductance microscopy (SICM) relies on the ionic current flowing through a nanocapillary in electrolyte. The ionic current strongly depends on the presence of a surface in the vicinity of the probe tip, hence the nanocapillary can be used as a nanoscale proximity sensor, without ever touching the sample. The sensing distance where the current drops is dependent on the pipette’s pore diameter^25^. SICM is therefore suitable for sensitive biological systems such as neurons, even with high-aspect ratio topographies with the invention of the hopping mode scanning modality^24^.

The use of combined SICM and stimulated emission depletion microscopy for topographical imaging was demonstrated to provide additional information about the structure of cytoskeletal components^26^, but lacked the ability to perform the complementary correlative recording on the same setup. Here we present a combined SICM-SOFI approach for in-situ, correlative single-cell imaging.

## Results

Our pipeline for a combined SICM/SOFI approach is based on correlative imaging with a home-built high speed SICM setup (Fig. 1a), operating in hopping mode^24^, combined with a SOFI workflow for either two- or three-dimensional sample imaging. SOFI increases the resolution of the widefield microscope by using the statistical information from stochastic intensity fluctuations and calculating the cumulants of the intensities between the individual pixels and image planes for resolution improvement in 2D and 3D respectively^11^, (Fig. 1b). The lateral resolution of SICM is approximately equal to three times the radius of the pipette^25^. In this work, we exclusively use nanocapillaries with a radius of 40-60 nm, which results in a lateral resolution comparable to SOFI. For increased resolution, glass nanocapillaries smaller than 10 nm radius can be fabricated^27^. Such nanocapillaries are however highly susceptible to clogging and are therefore not suitable for long-term measurements.

**Figure 1.**
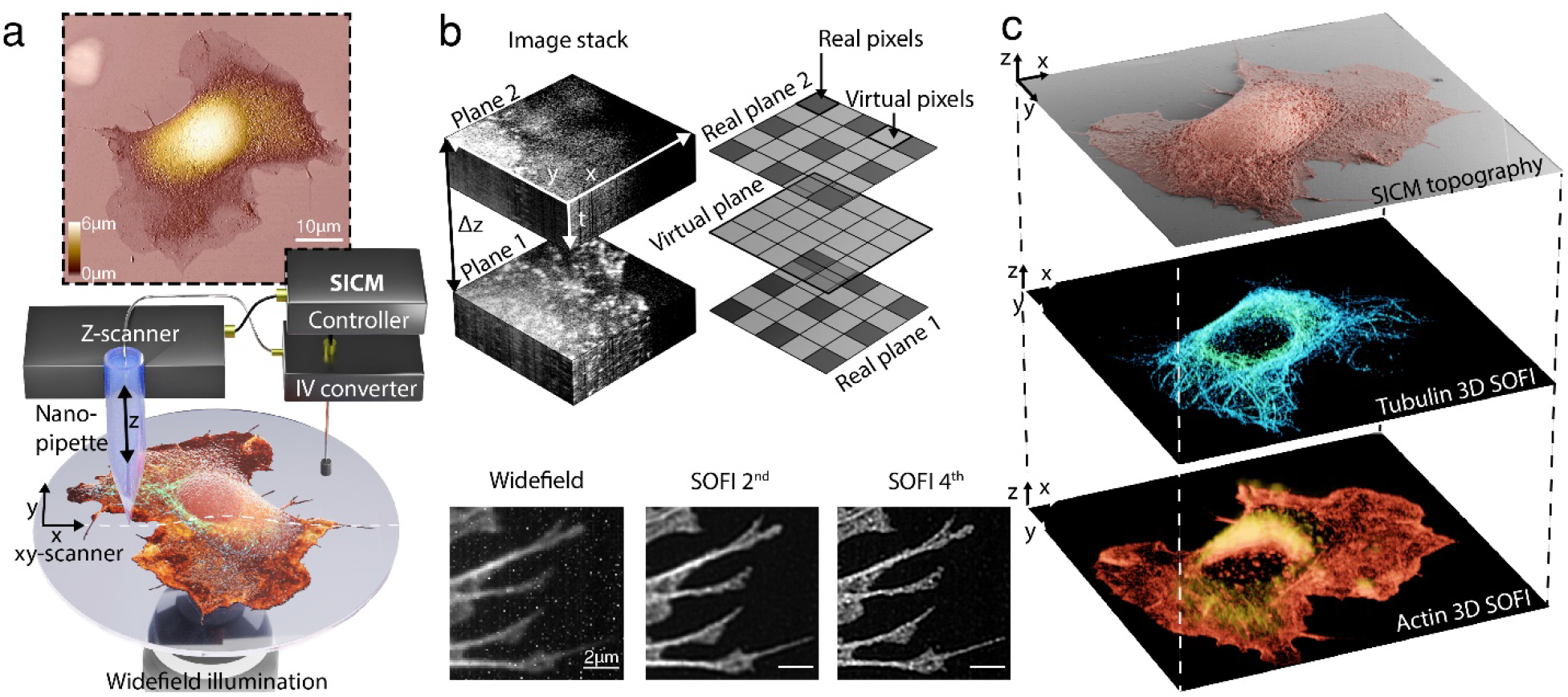
Combined scanning ion conductance microscopy (SICM) and super-resolution optical fluctuation imaging (SOFI) for three-dimensional topography-fluorescence correlative imaging. **(a)** Schematic illustration of the SICM-SOFI setup. In SICM, a current is generated through a nanopipette and modulated as a function of the z-position. The image generated reflects a three-dimensional topographical view of the surface at high lateral and axial resolution (top panel). The lateral resolution in SICM ranges from 150 nm to 30 nm and the axial resolution from 5 nm to the sub-nanometer range. Since the scanning head is mounted on top of the widefield setup, fluorescence imaging can be performed simultaneously. **(b)** Conceptual visualization of the SOFI principle. The recorded image stack, containing time traces of independently fluctuating fluorophores is used for analysis. A SOFI image is generated by computing cross-cumulants, creating virtual pixels between adjacent real pixels. The cross-cumulant computation principle can be applied in the axial direction as well if multiple sample planes are simultaneously acquired. For the n^th^ order SOFI image, the cumulant PSF volume is raised to n^th^ power thus giving the resolution improvement of a factor of 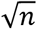. The inset images of the actin structure at the bottom of the **(b)** panel represent a standard deviation of the image stack (widefield), 2^nd^ and 4^th^ order of linearized SOFI images with corresponding lateral resolution values estimated with an image decorrelation-analysis algorithm of 420 nm, 194 nm, and 93 nm, respectively. **(c)** Schematics of correlative SICM and two-color 3D SOFI imaging.

To get label-specific information about the cytoskeletal structure of the cells, we performed SOFI with both traditional and self-blinking fluorescent dyes. Self-blinking dyes enable high-order SOFI analysis, that can be applied to various densities of temporally fluctuating fluorophores in order to gain spatial resolution and improve optical sectioning^14^. SOFI has the advantage of tolerating higher labelling densities^12^ compared to single-molecule localization microscopy, thus decreasing the acquisition time required to the order of 10 s of seconds per SOFI frame^28^. Subsequently, we correlated SICM data with two-color 2D and 3D SOFI images, that provided high-resolution single-cell maps showing a distribution of the actin and tubulin cytoskeletal proteins (Fig. 1c).

Before combining the two techniques within one instrument, we tested the 2D SOFI approach with self-blinking dyes on a dedicated setup (Fig. 2 a-b). We achieved up to 72 ± 3 nm resolution for 4^th^ order SOFI images for microtubules labelled with the commercially available Abberior FLIP-565 dye and 101 ± 9 nm for actin labelled with custom synthesized f-HM-SiR^29^, compared to a widefield resolution of 437 ± 106 nm and 480 ± 30 nm (mean ± s.d., N=8 images for SOFI 2D resolution measurements) respectively. Resolution was estimated using an image decorrelation analysis algorithm^30^, which allowed us to estimate the resolution image-wise. The calculated values agreed with theoretical values expected from SOFI analysis (Supplementary Fig. 1). Low-intensity (275 W/cm^2^ of 632 nm, 680 W/cm^2^ for 561 nm) illumination was used, resulting in minimal bleaching and long bleaching lifetimes of 406 ± 168 s for Abberior FLIP-565 and 625 ± 130 s for f-HM-SiR (mean ± s.d, N=8 images for SOFI 2D bleaching measurements) (Supplementary Fig. 2). Labelling density for both dyes was optimized for SOFI imaging, however both datasets were also processed with SMLM software^31^, which revealed that SOFI analysis was comparable in terms of resolution to the SMLM approach in low SNR conditions. Furthermore, analysis showed artefacts in high-density regions, non-optimal for SMLM analysis. The resolution, estimated with an image decorrelation analysis algorithm, was found to be 56 nm for tubulin and 98 nm for actin (Supplementary Fig. 3). The excellent optical sectioning and contrast provided by SOFI allowed to routinely resolve individual actin filaments (Fig. 2a, Supplementary Fig. 4). Subsequently, the corresponding cells were imaged in a custom SICM using an adaptive hopping mode. This yielded the topographical map of the cell membrane (Fig. 2c) resolving individual microvilli and filopodia membrane structures. Fluorescence and SICM images were registered based on the features from the topography map and the SOFI actin channel (Fig. 2d). Filamentous actin (f-actin) is known to be located in the lower part of the cell volume and correlates well with the topography of the cell boundary^32^ in COS-7 cells, that allowed us to simplify the registration process (Supplementary Fig. 5).

**Figure 2.**
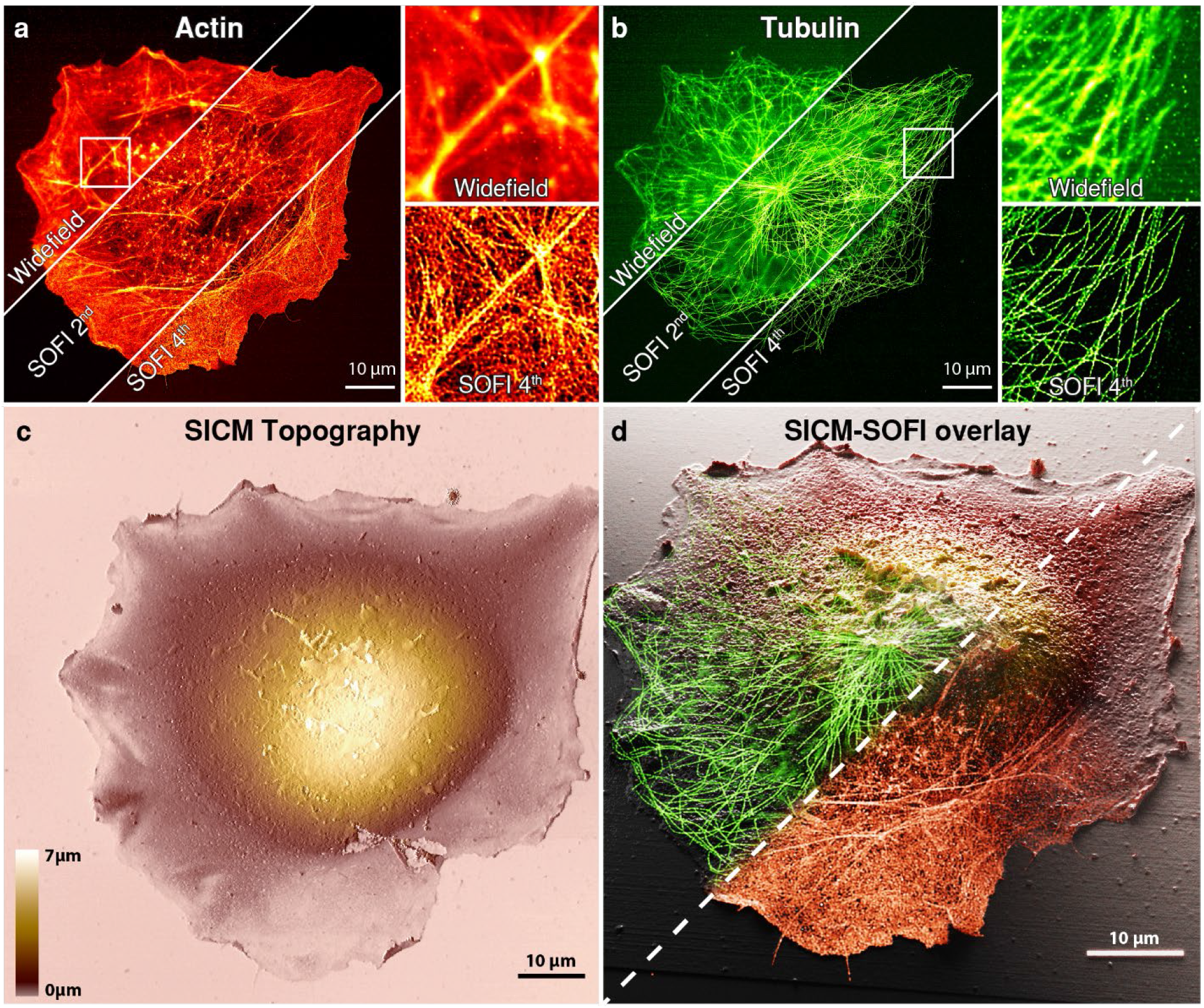
Correlative 2D SICM-SOFI imaging cytoskeletal components of a fixed COS-7 cell. **(a)** Large field of view (100 x 100 μm) 2D SOFI imaging of filamentous actin labeled with a phalloidin-f-HM-SiR self-blinking dye conjugate. **(b)** Subsequently, immunostained tubulin labelled with the self-blinking Abberior FLIP-565 dye was imaged. 10 x 10 μm zoom-ins of the standard deviation of the image stacks (widefield) and 4^th^ order SOFI images are shown for **(a)** and **(b)** panels. 635 nm flat-fielded laser excitation was used for imaging f-HM-SiR and a 561 nm laser line for Abberior FLIP-565. Imaging was performed in a 25% glycerol and PBS mixture at pH = 8. **(c)** Corresponding topographical SICM image of the same cell. **(d)** 3D rendering of a correlative SICM-SOFI overlay. The tubulin and actin channels are shown as a 2D plane, while topographical SICM information is used for the height representation in the Blender 3D software.

SOFI is not limited in providing the resolution improvement laterally, but it can also be used to improve axial resolution and sectioning by using information from multiple image planes acquired simultaneously. To demonstrate 3D SOFI’s capability of volumetric single cell imaging, we performed two-color 3D SOFI imaging for the same cytoskeleton components with a multiplane SOFI approach based on an image splitting prism^33^ (Fig. 3a-b), allowing us to calculate cumulants in 3D. However, the trade-off for 3D imaging capability is a reduced signal-to-noise ratio, which makes it difficult to perform high-order SOFI analysis, therefore we only show up to 3^rd^ order 3D SOFI images of microtubules labelled with the Abberior FLIP-565 dye and actin labelled with Alexa-647. (Supplementary Fig. 6). The acquired 8 physical planes resulted in 22 total planes spaced 116 nm apart after 3^rd^ order SOFI computation, giving a 3D volume of 50 x 60 x 2.45 μm with a lateral resolution comparable to the 3^rd^ order 2D SOFI results (Supplementary Fig. 1) A single color 3D SOFI image required as little as 3 min to acquire. This facilitates the screening of a large number of 3D cell volumes (Supplementary Fig. 7).

**Figure 3.**
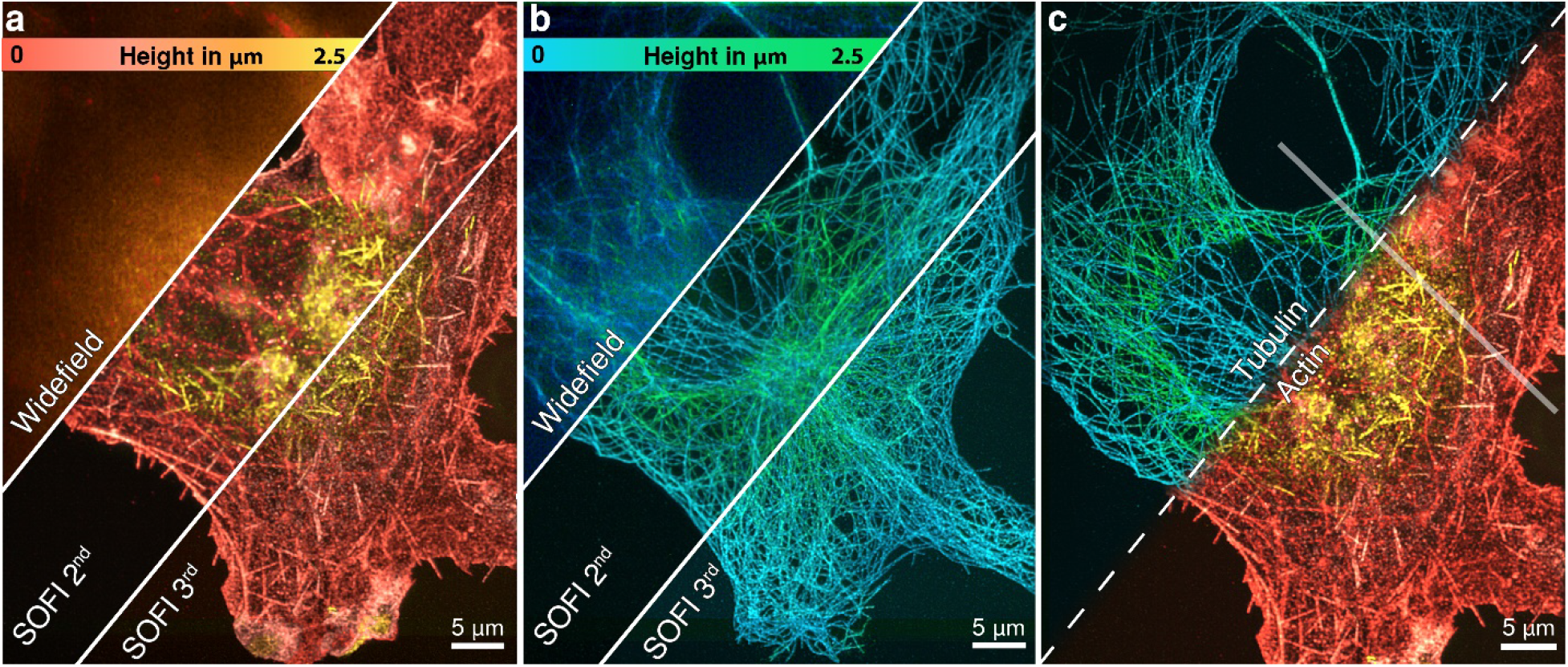
Multi-plane 3D SOFI imaging of cytoskeletal components in a fixed COS-7 cell. **(a)** 3D SOFI imaging of filamentous actin labeled with phalloidin conjugated to Alexa-647 and tubulin **(b)** labelled with the Abberior FLIP-565 self-blinking dye. A sample volume of 2.45 x 65 x 55 μm was recorded with 8 equally spaced physical planes, resulting in 22 image planes after 3^rd^ order SOFI processing. Flat-fielded 635nm and 532 nm laser lines were used for the excitation. Alexa-647 dye was imaged in ROXS imaging buffer with 10 mM MEA, while the Abberior FLIP-565 dye was imaged in a 50% glycerol and PBS solution at pH = 7.5. Height is represented in color scales displayed for each channel. **(c)** Co-registered 3D 3^rd^ order SOFI volumes of tubulin and filamentous actin. Co-registration was performed based on brightfield microscopy images acquired after recording each fluorescence channel. Semi-transparent line shows the cross-sections displayed in **Fig. 5 (a-b)**.

We then correlated the 3D SOFI information with the 3D SICM images, which have an order of magnitude higher axial resolution. The samples were moved to the dedicated SICM setup (Fig. 4a) to resolve microvilli at the surface of the cell (Fig. 4b-c). Finally, we correlated SICM and 3D SOFI information (Fig. 4d), which resulted in a detailed picture of the cell volume showing actin colocalized with microvilli protrusions over the whole cell surface (Fig. 4e).

**Figure 4.**
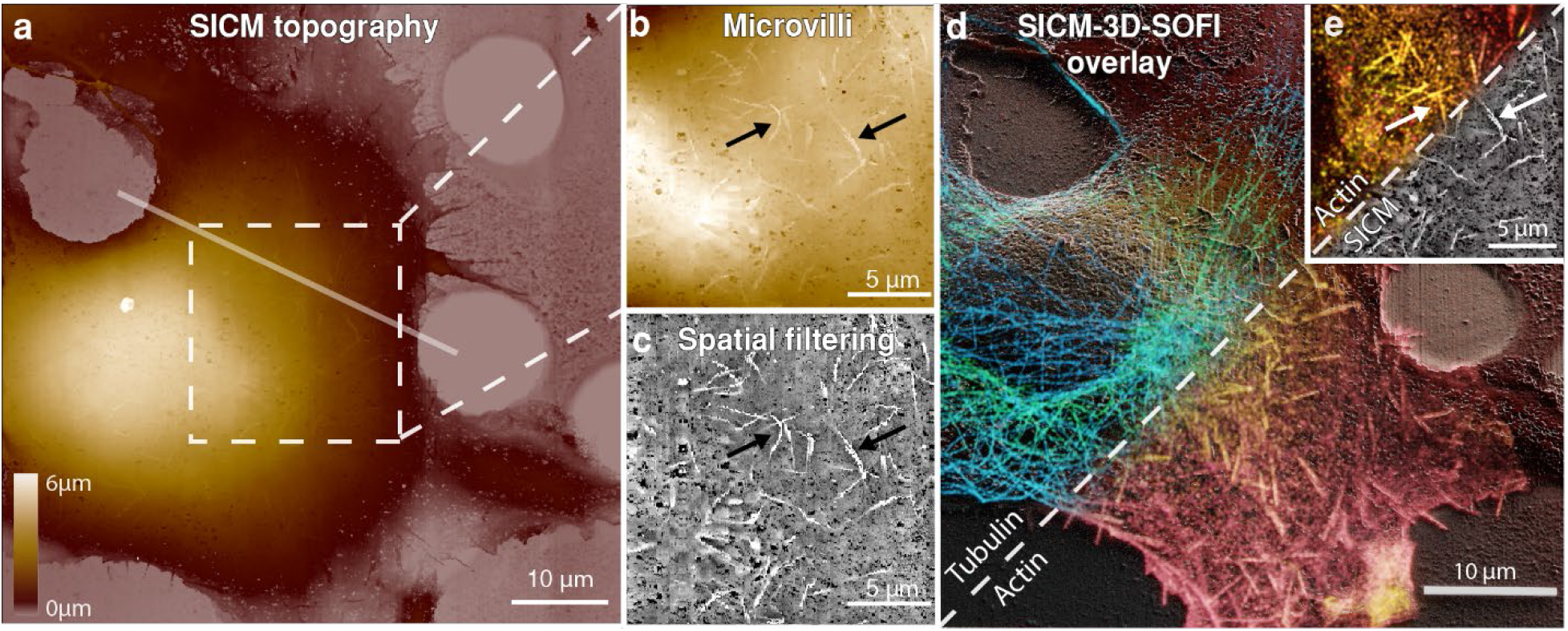
Correlative SICM and 3D SOFI imaging of cytoskeletal components in a fixed COS-7 cell. **(a)** Topographical SICM map of a fixed COS-7 cell imaged after two-color 3D-SOFI acquisition. Corresponding fluorescence images are shown in **Fig 3**. The scan resolution is 1024 x 1024 pixels over an 80 x 80 μm area with a corresponding pixel size of 78 nm. The pixel acquisition rate was 200 Hz with a hopping height of 6 μm. Semi-transparent line shows the cross-section area displayed in Fig. 5 **(a-b)**. **(b)** Leveled zoom-in of the upper part of the cell by mean plane subtraction, showing the topography of microvilli (marked with black arrows) on the surface of the cell. **(c)** Spatial band-passed filtered SICM image to highlight multiple microvilli structures. **(d)** 3D rendered correlative SICM and two-color 3D-SOFI overlay. The tubulin and actin channels are rendered as volumes consisting of 22 planes, while topographical SICM information is used for a height representation in the Blender 3D software (Supplementary movie 1). **(e)** Correlative SOFI and SICM overlay (white arrows correspond to the black arrows in **(b-c)**).

We have further interpreted the correlative information and compared the topographical maps of multiple cellular structures with two fluorescent channels which we acquired. Taking advantage of a multiplane SOFI approach, we compared the SICM topography with volumetric localization of actin (Fig. 5a) and tubulin (Fig. 5b) in a vertical section, which is marked in Fig. 3c and Fig. 4a. Patches of actin are distributed in the top and bottom parts of the cell, while tubulin is distributed homogenously within the cell volume. We also measured the correlation values for different cellular structures such as filopodia (Fig. 5c), microvilli (Fig. 5d) and microtubules (Fig. 5e) and calculated Pearson correlation coefficients between the pairs of actin, tubulin and SICM topography channels. The physical access to microtubules by SICM was obtained by partially removing the top membrane using Triton X-100 detergent during the permeabilization step^34^ (Supplementary Fig. 8). We found high correlations between the SICM/actin channels for filopodia and microvilli structures and for SICM/tubulin channels (Fig. 5f) for the chemically unroofed cell.

**Figure 5.**
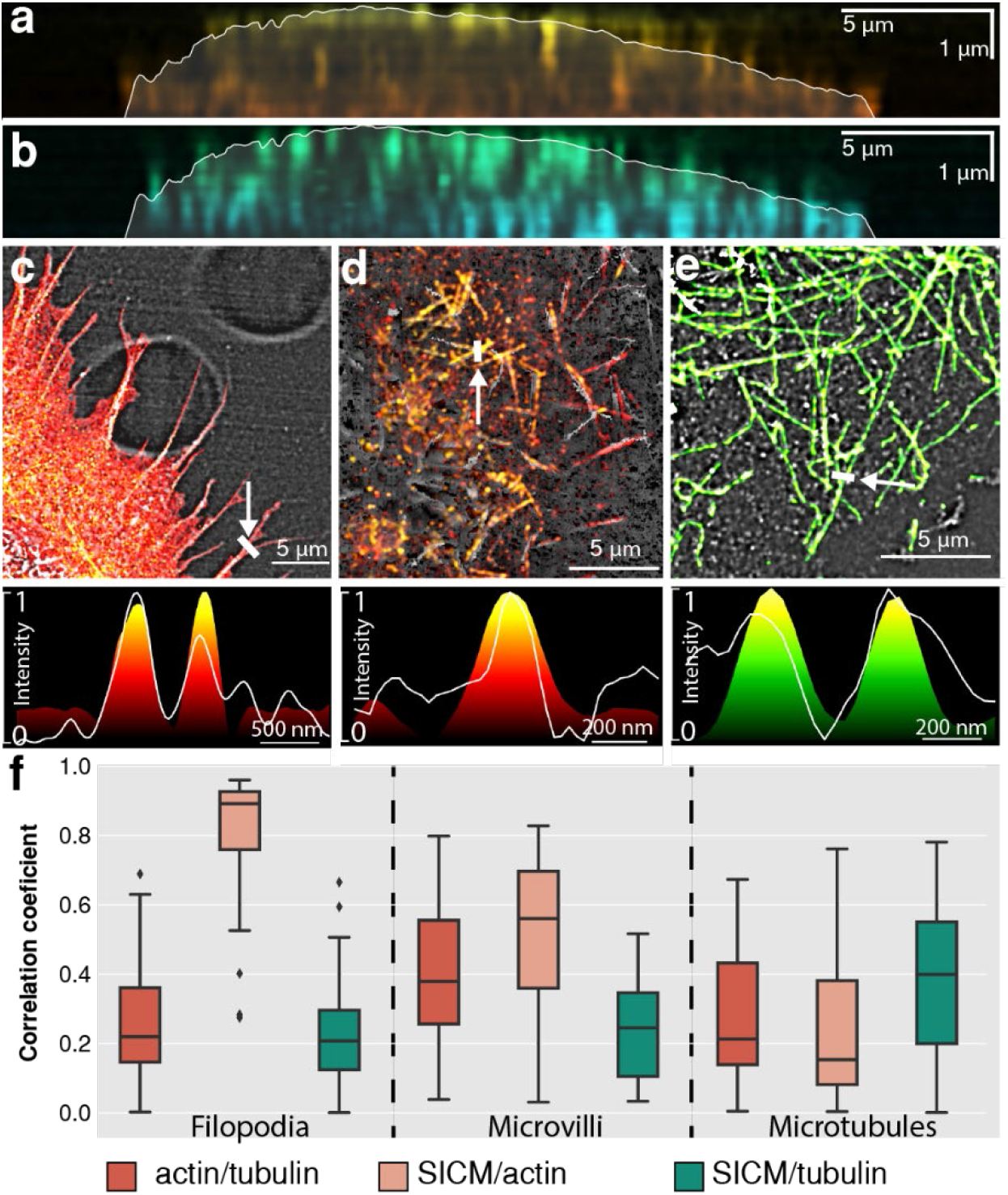
Volumetric distribution of cytoskeletal components from correlative SICM-3D-SOFI images. Vertical cross-sections of actin **(a)** and tubulin **(b)** in a fixed COS-7 cell marked in **Fig. 3c** and **Fig. 4a** overlaid with the SICM topography displayed as a semi-transparent contour. **(c-e)** Correlative imaging of different cellular components resolved by SICM: filopodia overlaid with fluorescently labeled f-actin from 3D-SOFI **(c)**, microvilli overlaid with fluorescently labeled f-actin from 3D-SOFI **(d)** and microtubules of chemically unroofed COS-7 cells overlaid with fluorescently labeled tubulin from 3D-SOFI **(e)**. Normalized intensity profiles are displayed below each image. For overlays only, the intensity values of SOFI images were projected on spatially filtered SICM images for a better structural representation. **(f)** Normalized Pearson cross-correlation coefficients of normalized height and intensity cross-sections for both SICM topography and fluorescence channels. Cross-sections (N=30) were manually selected (Supplementary Fig. 11) from the SICM image for each of the features. Correlation coefficients were measured on filipodia structures, microvilli structures and microtubules. Significantly higher (p < 0.05, two-sided t-test) correlation values were identified for filopodia (SICM)/actin, microvilli (SICM)/actin (3D-SOFI) and microtubules (SICM)/tubulin (3D-SOFI).

The establishment of a correlative SOFI/SICM pipeline on different instruments paved the way for a combined instrument. For correlative live-cell imaging, we combined the prototype SICM instrument with a 2D SOFI capable widefield microscope. We performed a correlative measurement of cytoskeletal proteins and cell morphology revealing the correlated dynamics of actin and membrane topography at the subcellular level. We transfected COS-7 cells for cytoskeletal proteins of actin filaments (actinin or actin) fused with mEOS-2 photo-switching fluorescent proteins in order to achieve stochastic fluctuations in the fluorescence signal. Cells were then scanned in a consecutive manner with SICM and imaged with SOFI in a custom-design chamber with an environmental control. Recordings of 300 frame-long stacks of the fluorescence signal were performed after each SICM frame, resulting in a combined SICM/SOFI acquisition time of 3 min/frame for images of parts of the cell, and 10 min for whole cell imaging (Supplementary Figs 9–10). The 2^nd^ order SOFI images in Fig. 6a show the actin filaments with a 173 ± 20 nm (mean ± s.d, N=6 images for live-cell SOFI 2D resolution measurements) lateral resolution. After recording, SICM and SOFI images were aligned, revealing correlated dynamics of cytoskeletal proteins and membrane topography (Fig. 6b-c). During the SICM imaging, there was no laser applied, resulting in two-minute dark intervals between SOFI frames. This significantly reduced the photodamage enabling us to record time-lapse sequences of 10-15 SOFI frames for up to 42 min without observable cell negative phototoxic effects (Supplementary Fig. 9).

**Figure 6.**
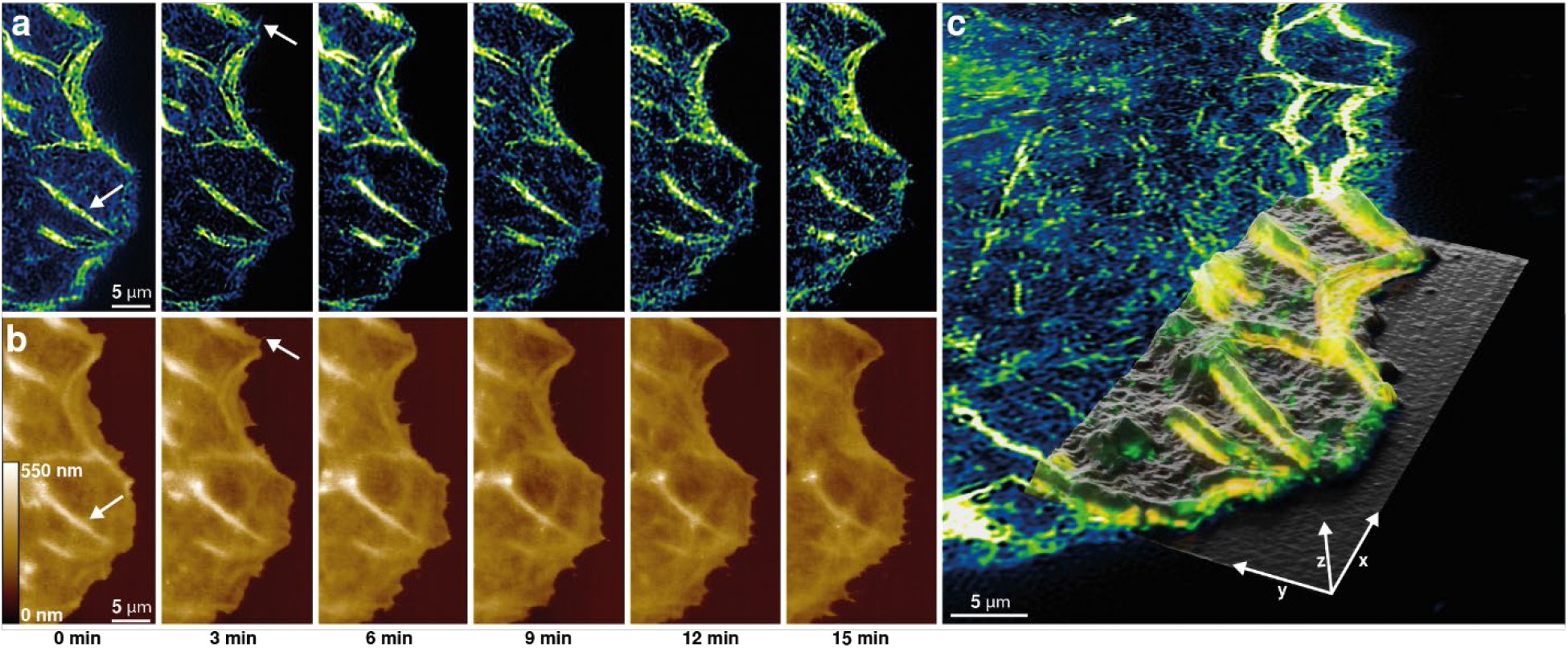
Live-cell SICM/SOFI imaging of cytoskeletal actinin dynamics of COS-7 cell. **(a)** 2^nd^ order SOFI images of actinin-mEOS-2 in transfected COS-7 cells reconstructed from 250 frames each, thus allowing to achieve 12.5 s temporal resolution per SOFI frame. **(b)** Corresponding SICM height maps recorded at 150 s temporal resolution per SICM frame. **(c)** 3D render of SICM height map aligned to the full SOFI image revealing the correlative dynamics of actinin in a live-cell (Supplementary movie 2).

## Discussion

We have established correlative membrane topography and cytoskeleton imaging provided by combined SICM and SOFI modalities for fixed COS-7 cells both in 2D and 3D on separate imaging setups preserving the state-of-the-art capabilities of both imaging modalities. For imaging densely-labelled samples we used novel self-blinking dyes suitable for high-order SOFI imaging. Additionally, the 3D SOFI approach allowed us to retrieve information about the cytoskeletal protein distribution within the cell volume. Subsequently acquired SICM high-resolution axial topography provided detailed volumetric cell-mapping. Finally, we have performed simultaneous SICM/SOFI live-cell imaging in a combined setup for routinely obtaining correlative measurements in vitro.

Previous studies have demonstrated that a combination of topographical and biochemical sample information is a powerful tool which can provide a comprehensive picture of cellular activity. Correlative SPM and super-resolution microscopy studies, involving techniques such as SMLM^35^, stimulated emission depletion (STED)^26,36^, structured light illumination (SIM)^21^ have been demonstrated. SOFI, as a computational method, does not require a complex optical setup design compared to SIM or STED approaches. We show that it can be easily implemented on existing SPM/SICM setups with only few optical components required. Simultaneous SICM measurements can be used to directly detect the cell topography changes related to apoptosis^37^ thus acting as a detector for phototoxicity. Furthermore SICM has been shown to outperform AFM in live-cell imaging^22^ due to its non-contact nature, which is crucial in imaging sensitive samples. On the other hand, AFM has demonstrated its superiority over SICM in terms of imaging resolution^38^. AFM is also more versatile in terms of retrieving information about the nanomechanical sample properties^39^ and can even be used for molecular-specific imaging^40^. Due to the fragile nature of the glass nanocapillary, SICM is less forgiving of imaging mistakes that can lead to fracture of the capillary. This is exacerbated by the fact that no pre-fabricated SICM capillaries are commercially available.

We demonstrated the ability of the combined method for biological studies of the dynamic cell morphology and cytoskeletal architecture inside the cells. This could be used to investigate open questions in membrane trafficking^3^, cell migration^41^ or infection. Due to the electrical nature of SICM measurements, it can also retrieve surface charge information^42^, thus adding a whole new dimension for the measurement. The ability to additionally map the charge of a cell membrane is particularly relevant for problems such as clustering of membrane proteins^43^, membrane curvature influence on density of membrane proteins and lipids^44^, lipid-rafts^45^, behaviour of voltage-gated ion channels^46^, etc. While there are some remaining technological challenges in terms of usability, the combination of the multimodal SICM and flexible SOFI has the potential to become a routine live-cell imaging modality capable of tackling challenging biological problems.

## Methods

### 2D widefield fluorescence imaging setup

A home-built widefield microscopy setup described previously^14^ was used (Supplementary Fig. 12). The setup has four laser lines for illumination: a 200 mW 405 nm laser (MLL-III-405-200mW, Roithner Lasertechnik), a 1 W 632 nm laser (SD-635-HS-1W, Roithner Lasertechnik), a 350 mW 561 nm laser (Gem561, Laser Quantum) and a 200 mW 488 nm laser (iBEAM-SMART-488-S-HP, Toptica Photonics). The beam of the 635 nm laser is flat fielded by coupling it into the multimode fiber and passed through a speckle reducer (Optotune, LSR-3005-17S-VIS) similarly to that described previously^47^. All laser lines are collimated, expanded and focused in the back focal plane of the water immersion 60x objective (Nikon SR Plan Apo IR 60× 1.27 NA WI) resulting in the effective FOV of 100×100 μm with a pixel size of 107 nm. Fluorescence signal is then filtered using a quad-line dichroic mirror (ZET405/488/561/640, AHF Analysetechnik) and emission filter (R405/488/561/635 flat, AHF Analysetechnik). Additional band-pass emission filters (596/83 or 685/70, Chroma) were used for sequential two-color imaging. Finally, the light is focused on a sCMOS camera (ORCA Flash 4.0, Hamamatsu; back projected pixel size of 108 nm). Z stabilization is achieved with a PID controller using a total internal reflection from a coverslip using 780 nm laser diode [model] reflecting from a sample at the critical angle. Axial positioning is achieved with a nano-positioning stage (Nano-Drive, MadCityLabs) using a custom-written software in LabVIEW environment. Lateral sample position is controlled by a Scan-plus IM 120×80 (Marzheuser) stage. Image sequence acquisition is done in Micromanager software.

### 3D widefield imaging setup

Multiplane SOFI imaging was performed on a home-built widefield microscope (Supplementary Fig. 13) with a simultaneous 8-plane detection system using an image splitting prism^48^. Four laser lines were used for illumination: a 120 mW 405 nm laser (iBeam smart, Toptica), an 800 mW 635 nm laser (MLL-III-635, Roithner Lasertechnik), 200mW 488nm laser (iBEAM-SMART-488-S-HP, Toptica Photonics) and an 800 mW 532 nm laser (MLL-FN-532, Roithner Lasertechnik). The beam of the 635 nm laser is flat fielded by coupling it into the multimode fiber. All laser lines are collimated, expanded and focused in the back focal plane of the water immersion 60x objective (Olympus UPLSAPO 60XW 1.2 NA). The fluorescence signal is then filtered using a dichroic mirror (zt405/488/532/640/730rpc, Chroma) and quad band emission filter (405/488/532/640m Chroma). Additional band-pass emission filters (582/75 or 685/70, Chroma) were used for sequential two-color imaging. An image-splitting prism was placed behind the last lens and splits the signal into 8 images recorded by two synchronized sCMOS cameras (ORCA Flash 4.0, Hamamatsu; back projected pixel size of 111 nm). Each image plane is equally spaced 350 nm apart, resulting in a total imaging volume of 50 × 60 × 2.45 *μm*^3^. The sample is positioned in XYZ by using a hybrid piezo nanopositioning stage (3-PT-60-F2,5/5) and Motion-Commander-Piezo controller (Nanos Instruments GmbH). Synchronization of cameras, imaging acquisition and general setup control is done with a custom-written software in LabVIEW environment.

### Scanning ion-conductance microscopy setup

Scanning probe microscopy was performed with a custom-made scanning ion conductance microscope. The sample was actuated in X and Y by a piezo-stage (piezosystem Jena TRITOR102SG). The nanocapillary was moved in Z by a home-built actuator, operated in hopping mode. Borosilicate and quartz nanopipettes were fabricated with a CO-2 laser puller (Model P-2000, Sutter Instruments) with a radius from 40 nm to 60 nm and 20 nm to 40 nm respectively.

### Combined 2D widefield fluorescence /SICM setup

A home-built widefield setup is assembled in combination with a SICM scanner (For more detail see SICM setup) mounted atop an inverted Olympus IX071 microscope body (Supplementary Fig. 14). For sample excitation, a four-color (405 nm, 488 nm 561 nm, 647 nm) pigtailed Monolithic Laser Combiner (400B, Agilent Technologies) is used. Light is collimated and focused to the back focal plane of the oil-immersion high-NA objective (Olympus TIRFM 100x, 1.45 NA) by using a custom built TIRF illuminator. The fluorescence signal is then filtered using a dichroic mirror (493/574 nm BrightLine, Semrock) and a band emission filter (405/488/568 nm stop line, Semrock). Finally, the light is focused on an sCMOS camera (Photometrics, Prime 95B; back projected pixel size of 111 nm). Coarse lateral sample positioning is done with a mechanical stage, while fine positioning is achieved with a SICM XY piezo scanner (Piezosystemjena TRITOR102SG). Image stacks for SOFI are recorded with a Micromanager software, while laser control and sample positioning are achieved in custom-written LabVIEW software.

### Coverslip fabrication

High precision No. 1.5 borosilicate 25 mm coverslips (Marienfeld) were patterned with a custom layout (Supplementary Fig. 15) by using a commercial UV excimer laser patterning setup (PTEC LSV3). It allowed to create a user-friendly sample map, which was crucial for further SOFI-SICM correlation experiments while transferring the sample between different setups. After patterning, coverslips were cleaned with piranha solution, washed in MiliQ water, dried with N_2_ flow and kept dry for further use. Before use, coverslips were cleaned with oxygen plasma cleaner (Femto A, Diener electronic GmbH) for 660 s at maximum power setting, washed with PBS (pH = 7.4) once and coated with 50 μM fibronectin solution in PBS (pH = 7.4) for 30 min at 37° C before seeding the cells.

### Cell culture

Cells were cultured at 37 °C and 5 % CO_2_. DMEM high glucose without phenol red medium (Gibco, Thermo Fisher Scientific) was used, containing 10 % fetal bovine serum (Gibco, Thermo Fisher Scientific), 1% penicillin-streptomycin (Gibco, Thermo Fisher Scientific) and 4 mM L-glutamine (Gibco, Thermo Fisher Scientific). Before seeding on coverslips, cells were detached from a flask with TrypLE (Gibco, Thermo Fisher Scientific) and 30 000 cells were seeded on 25 mm coverslips coated with fibronectin. Coverslips were washed twice with PBS before seeding the cells and 2 ml of DMEM medium was used. Cells were grown overnight (12-16 h) before fixation in 6-well plates.

### Sample fixation and staining

Cells were washed once with DMEM described previously and incubated for 90 s in a prewarmed microtubule extraction buffer, consisting of 80 mM PIPES, 7 mM MgCl_2_, 1 mM EDTA, 150 mM NaCl and 5 mM D-glucose with a pH adjusted to 6.8 using KOH with 0.3 % (v/v) Triton X-100 (AppliChem) and 0.25 % (v/v) EM-grade glutaraldehyde (Electron Microscopy Sciences). After 90 s the solution was exchanged to pre-warmed 4 % paraformaldehyde dissolved in PBS (pH=7.4) and samples were incubated for 10 min at room temperature. Afterwards, samples were washed thrice for 5 min with PBS on orbital shaker. Cells were kept for 5 min with a freshly prepared 10 mM NaBH4 solution in PBS on an orbital shaker in order to reduce background fluorescence. Step was followed by one quick wash in PBS, and two washes of 10 min in PBS on an orbital shaker. Samples were then additionally permeabilized to ensure antibody penetration with 0.1 % (v/v) Triton X-100 in PBS (pH=7.4) on an orbital shaker followed by additional wash with PBS. Finally, samples were blocked with freshly prepared blocking buffer consisting of 2 % (w/v) BSA, 10 mM glycine, 50 mM ammonium chloride NH4Cl in PBS (pH=7.4) for 60 min at room temperature or stored overnight at 4 °C for further staining. All chemicals were bought from Sigma Aldrich unless stated differently.

### Two-color sample staining

For incubation with antibodies and/or phalloidin, coverslips were placed on parafilm in a closed box in the dark, at high humidity to prevent coverslips from drying. 100 μL of staining solution was typically used for each coverslip. After blocking, samples were incubated with 2 % (v/v) primary anti-tubulin antibody (clone B-5-1-2, Sigma-Aldrich) in blocking buffer for 60 min at room temperature. Samples were washed with blocking buffer thrice for 5 min on orbital shaker. Coverslips were incubated with 2 % (v/v) secondary donkey anti-mouse-Abberior FLIP-565 antibody, which was labelled as described previously^14^. Samples then were kept in blocking buffer for 60 min and washed thrice for 5 min on orbital shaker. Samples were incubated for 10 min in 2 % (w/v) PFA in PBS (pH=7.4) as a post-fixation step followed by three 5 min washes with PBS on orbital shaker. After tubulin staining, actin was stained with 500 nM custom synthesized phalloidin-f-HM-SiR (Supplementary Fig. 16) or phalloidin-Alexa-647 (Sigma Aldrich) solutions in PBS by incubating for 60 min at room temperature. Samples were washed thrice for 5 min with PBS on orbital shaker and imaged immediately in imaging buffer.

### Imaging buffers

2D imaging with self-blinking dyes was performed in 50 % glycerol PBS solution at pH=8. The buffer was degassed with N_2_ flow for 30 min before use. 3D phalloidin-actin imaging was performed in an imaging buffer described previously^49^ containing 10 % (w/v) D-glucose, 20 % (v/v) glycerol, 50 mM TRIS, 10 mM NaCl, 2 mM COT, 10 mM MEA, 2.5 mM PCA and 50 nM PCD with a pH adjusted with HCl to 7.5. 3D imaging of tubulin labelled with an Abberior FLIP-565 was performed in 50 % (v/v) glycerol solution in PBS (pH=7.4). All imaging experiments were done in a sealed imaging chamber.

### Two-color SOFI imaging

2D SOFI imaging was performed with self-blinking dyes sequentially. Phalloidin-f-HM-SiR labelled actin was imaged first at 275W/cm^2^ 635 nm excitation, then 680 W/cm^2^ of 561 nm was used to image Abberior FLIP-565 labeled tubulin. The sample was kept on the setup for at least 30 min before imaging, to reduce thermomechanical drift during imaging. 16 000 – 30 000 frames were acquired for each channel for high-order SOFI analysis with a minimal photobleaching effects (Supplementary Figs. 2 and 17). Sample drift between two-color acquisitions was neglected for further analysis. 3D SOFI imaging was first performed with phalloidin-Alexa-647 labelled actin excited with a 632 nm laser at 3.4 kW/cm^2^ with a low power (5W/cm^2^) 405 nm activation laser to increase the population of fluorophores in a bright state. After imaging multiple cells, the oxygen-scavenging imaging buffer was changed to 50 % (v/v) glycerol in PBS for Abberior FLIP-565 labelled tubulin imaging. The sample was then imaged with the 532 nm laser at 3.5 kW/cm^2^. 4 000 frames for each image plane were recorded for high-order 3D-SOFI analysis. After recording each image stack, a subsequent brightfield-image was taken for a further alignment of 2-channel data. 50 ms exposure time was used for all imaging experiments. The data presented in the paper are from two distinct samples.

### SICM imaging and processing

The SICM imaging was performed in PBS (pH = 7.4). The hopping height ranged from 1 μm to 3 μm with hopping rates from 200 Hz to 1 kHz. The current setpoint used in the hopping actuation was 99% of the normalized current recorded. On fixed cells, SICM images with 1024×1024 pixels were generated after performing fluorescence imaging. Corresponding cells were found using markers described in a coverslip fabrication section. On live cells, SICM images with 512×256 pixels were generated in the combined SICM/SOFI system described previously. SICM images were further processed using Gwyddion^50^ and Fiji^51^.

### SOFI Image analysis and image alignment

2D and 3D SOFI image analysis was performed as described previously ^14,48^. Briefly, the image stack was drift-corrected using cross-correlation between SOFI sub-sequences before performing cumulant calculation. Different planes of the 3D image stack were registered based on acquired bead stack and images sequence was drift-corrected after the registration, assuming that the drift is homogenous within the volume. 4^th^ order SOFI images were used in 2D SOFI case, while 3^rd^ order SOFI images were taken for 3D SOFI. Images were further deconvolved with a Lucy-Richardson deconvolution algorithm using gaussian PSF model. Deconvolution settings were optimized using decorrelation analysis algorithm for resolution estimation ^30^. 2D and 3D SOFI analysis code was implemented in Matlab and is available upon request. After 2D and 3D SOFI processing, images were aligned with a topographical SICM map using custom-written Python code based on the hand-selected features in the actin channel and SICM image (**Supplementary Fig. 5**). For two-color 3D SOFI, two-color SOFI stacks were co-registered based on brightfield images and then aligned with SICM image based on the actin channel.

### Live-cell SOFI/SICM imaging

COS-7 cells were seeded according to the procedure described previously. After 12-16h cells were transfected with actinin-mEOS2 or Lifeact-mEOS2 (Supplementary information: Plasmid sequences) with a Lipofectamine 3000 (Thermo Fisher) according to the protocol provided by manufacturer. Cells were used for combined SICM-SOFI experiment 24 h after the transfection. Imaging was performed at 37 °C and 5 % CO_2_ in a custom-built imaging chamber in FluoroBrite DMEM media (Gibco, Thermo Fisher Scientific) in order to reduce autofluorescence. SICM and SOFI imaging was performed subsequently i.e. after each SICM image, a fluorescent stack of 300 frames with an exposure time of 50 ms was recorded. The first 50 frames were excluded from SOFI processing due to rapid intensity change upon 405 activation and only 250 frames were used for further SOFI analysis. Live-cell imaging was performed under the low-illumination intensity (500 W/cm^2^ of 561 imaging laser and 0.3 W/cm^2^ of 405 nm activation laser) in order to reduce phototoxicity.

### Correlative 3D SICM/SOFI data rendering

In order to fully expose the correlative 3D data, we have used the advanced open-source 3D rendering tool Blender 3D for data visualization. Normalized topographical SICM data was imported as a height map and scaled in the axial direction. Then, 2D SOFI data was overlaid by importing a co-registered fluorescence image and a custom-written shader was used for volumetric multiplane data rendering (Supplementary Fig. 18) to generate the final figures.

## Additional information (Supplementary Figures 1–18, plasmids used)

### Data availability

Raw and processed image data, together with scripts used for analysis are available on reasonable request from the corresponding authors.

## Acknowledgements

V.N. and A.R. acknowledges the support of the Max Planck-EPFL Center for Molecular Nanoscience and Technology and Swiss National Science Foundation through the National Centre of Competence in Research Bio-Inspired Materials. K.Y. and D.P.H. gratefully acknowledge funding by the Centre of Membrane Proteins and Receptors (COMPARE, Universities of Birmingham and Nottingham). GEF acknowledges the support by the European Research Council under grant number ERC-2017-CoG; InCell; Project number 773091. The authors thank Prof. Jens Anders f and his team from the University of Stuttgart for insightful discussions into high-speed transimpedance amplifiers.

## Author Contributions

V.N. and S.L. equally contributed to the paper by conceiving and refining general idea and writing the final manuscript with input from all the authors. V.N. prepared the cell samples, performed all optical imaging experiments and correlative SOFI image data analysis and the final data visualization. B.D and S.L. built the SICM setup, S.L. performed the SICM imaging and SICM data processing. Simultaneous SICM/SOFI imaging was performed by V.N. and S.L. K.G. contributed by creating and adapting labelling protocols. A.D. adapted and improved the existing SOFI analysis code. K.Y. and P.W. synthesized and provided f-HM-SiR-phalloidin conjugate. D.-P.H., R.W., A.R and G.E.F. supervised the work.

## Supplementary information

**Supplementary Figure 1.**
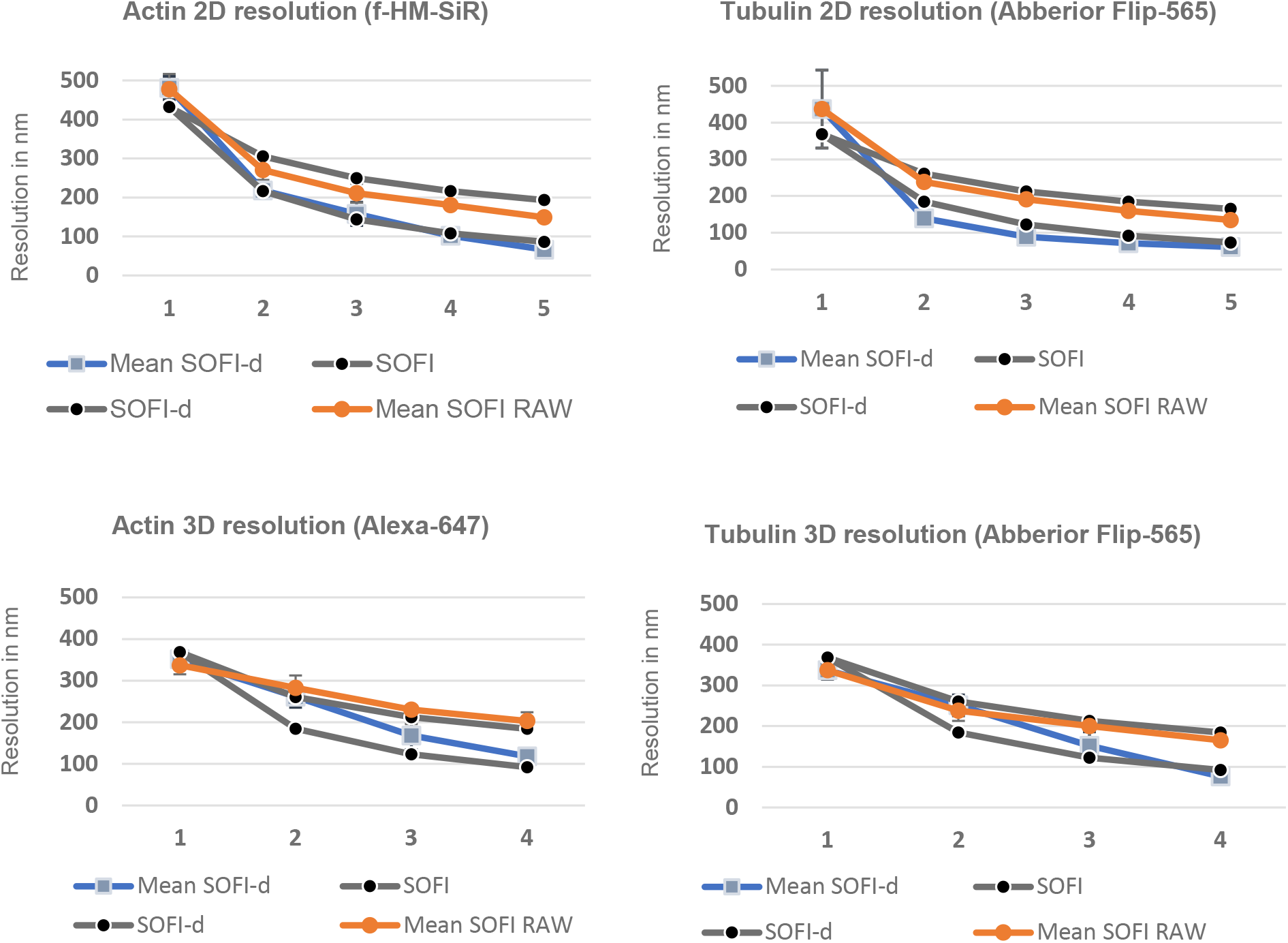
SOFI resolution estimation for two-color 2D and 3D SOFI images. Resolution was estimated with a parameter-free image resolution estimation algorithm ^3^. For 2D images, each value is calculated from 8 images. For 3D SOFI stack resolution was estimated in all planes (N=8, N=15, N=22, N=29) for corresponding SOFI orders. Theoretical values for SOFI and SOFI-d (deconvolved) based on the diffraction limited PSF size are also plotted.

**Supplementary Figure 2.**
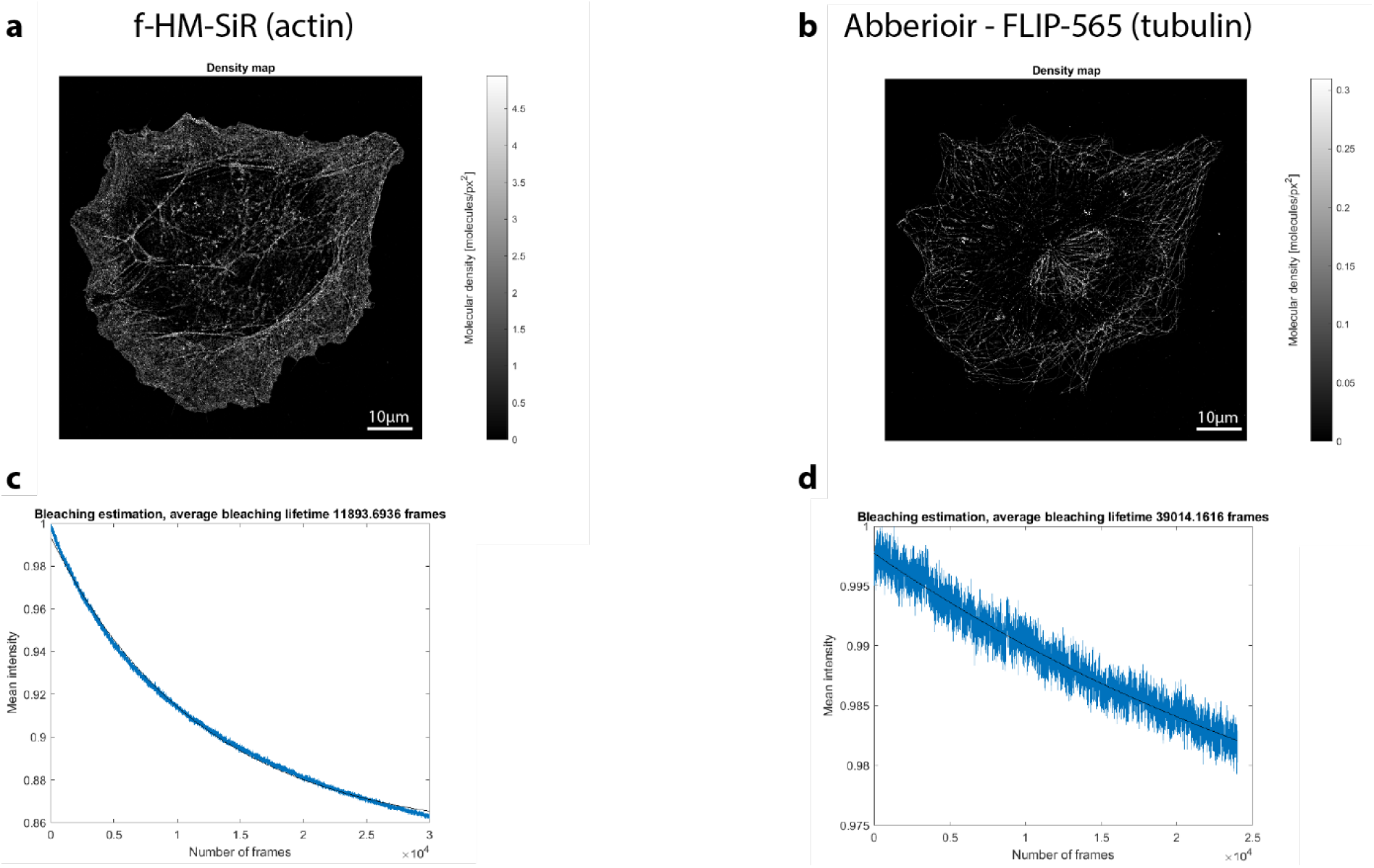
Labelling density and bleaching kinetics of self-blinking dyes. Molecular densities of f-HM-SiR (a) and Abberior-FLIP 565 (b) dyes from a representative image, together with bleaching curves estimated from the corresponding image stacks. Calculated average bleaching lifetimes (8 stacks with 30 000 frames) for f-HM-SiR dye was 406 ± 168 s and 625 ± 130 s for Abberior FLIP-565 (mean ± s.d, N=8 image stacks consisting of 30 000 frames).

**Supplementary Figure 3.**
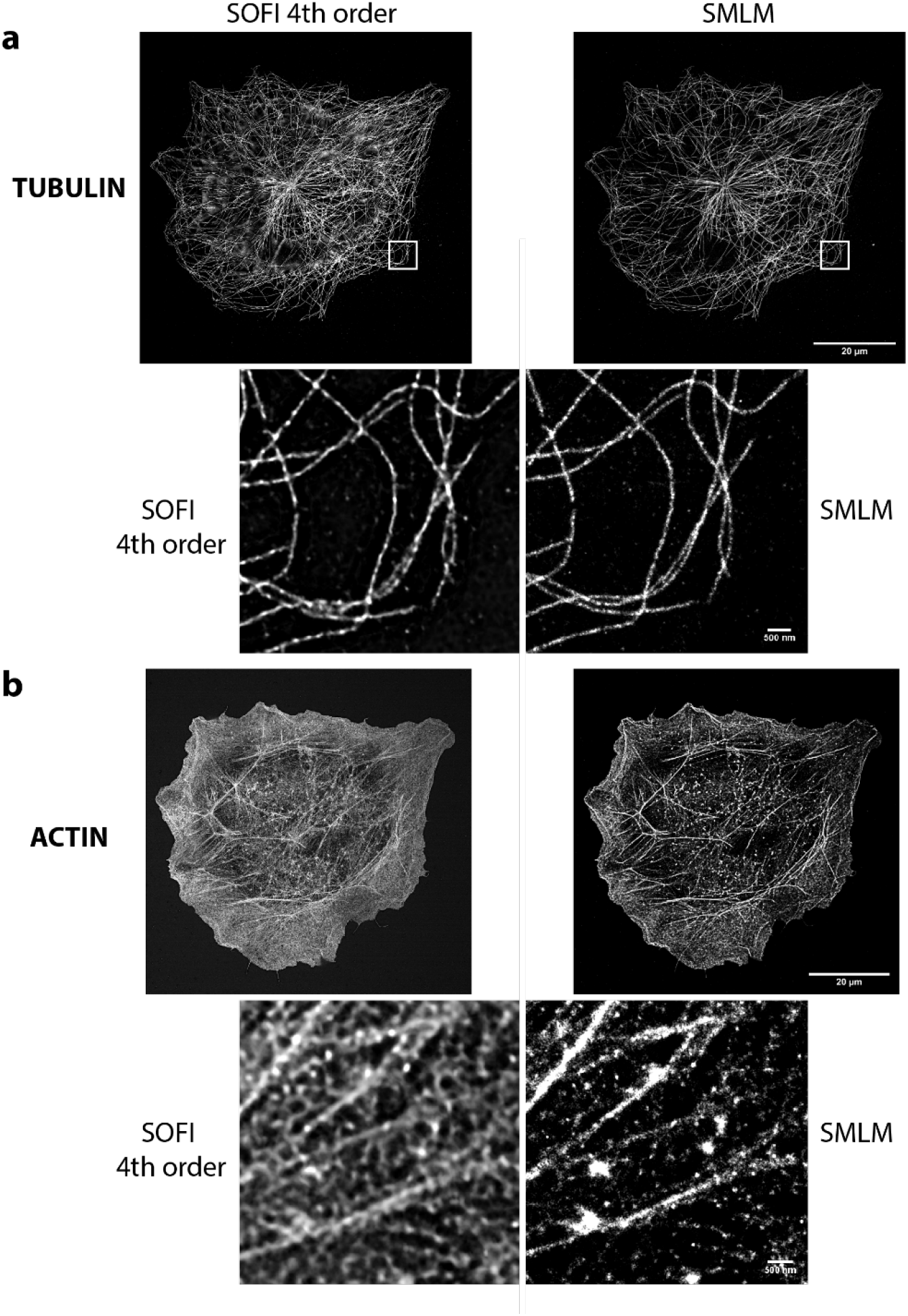
High order SOFI comparison with SMLM. a) 4^th^ order SOFI and SMLM (Thunderstorm) image quality comparison. Images stacks were processed with the latest version of Thunderstorm software^2^ by using single-emitter fitting function. It is visible that in sparse blinking conditions (Abberior FLIP-565 labelled microtubules) both approaches (SOFI and Thunderstorm) perform similarly, however for f-HM-SiR labelled f-actin SMLM seems to produce localization artifacts, that are expected for high-density data. Resolutions metric stated in the paper were computed with image decorrelation analysis algorithm^3^. The localization precision might be improved by using multi-emitter fitting or pre-processing tools such as HAWK^4^, however this is clearly outside of the scope for this study.

**Supplementary Figure 4.**
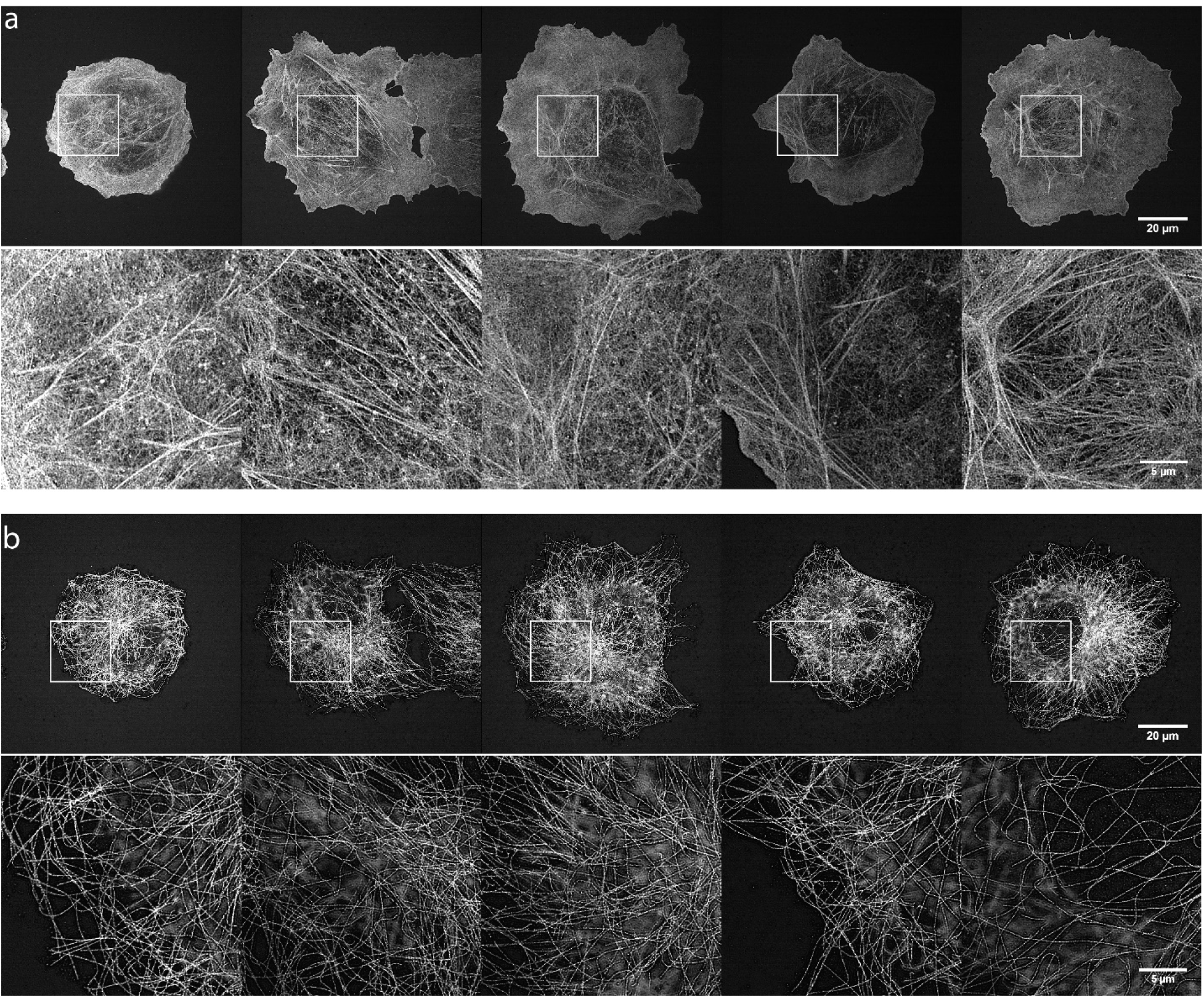
Multiple two-color 4^th^ order SOFI images. 4^th^ order SOFI images of phalloidin-f-HM-SiR labeled actin (a) and Abberior FLIP-565 immunostained tubulin (b). Zoom-ins of 25 μm are shown together with the whole 90×90 μm field of view images.

**Supplementary Figure 5.**
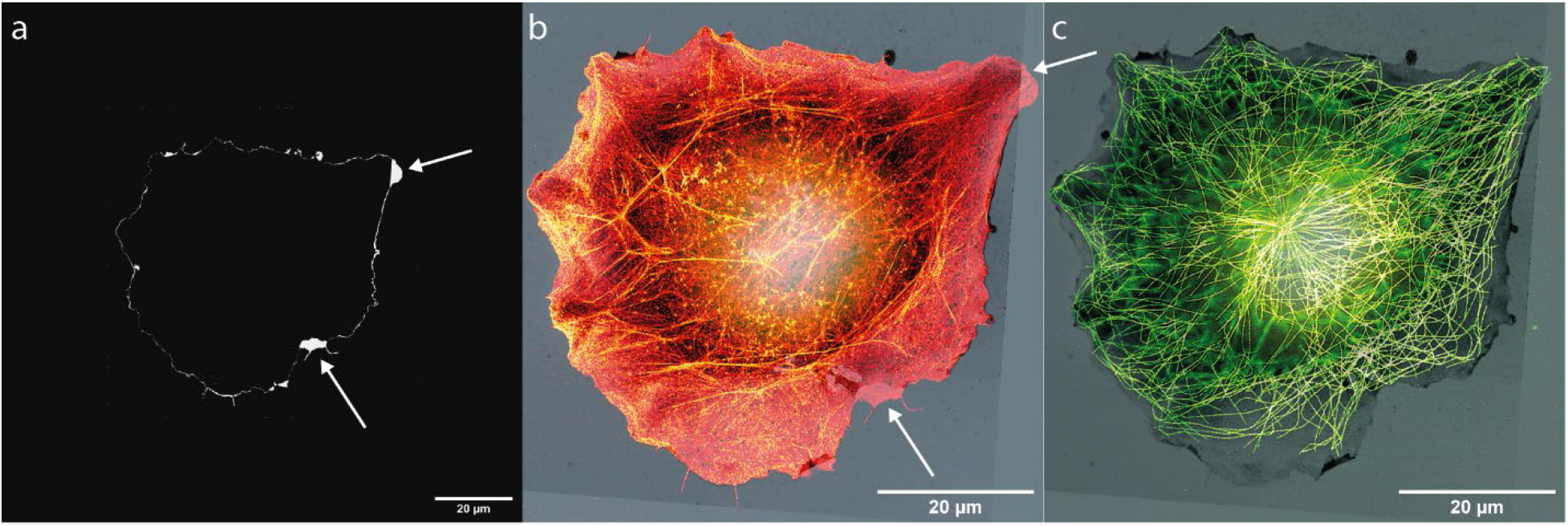
SICM SOFI image co-registration procedure. (a) Difference of the tresholded SICM and actin channels revealing the different cell features between SICM and actin channels. (b) SICM image overlayed with actin and (c) channels. Arrows are showing the ambiguities marked in image (a). 2D SOFI images were aligned with SICM topographical map based on the actin channel. Same features were manually depicted in both images and affine transformation matrix was computed using at least 10 corresponding marks. 2D tubulin image was transformed using the same transformation matrix assuming that the lateral drift between two SOFI images is neglectable. 3D SOFI image stacks were processed in the same way expect in this case two-color stacks were aligned based on the brightfield microscopy by phase correlation using the images recorded before acquiring each of the stack.

**Supplementary Figure 6.**
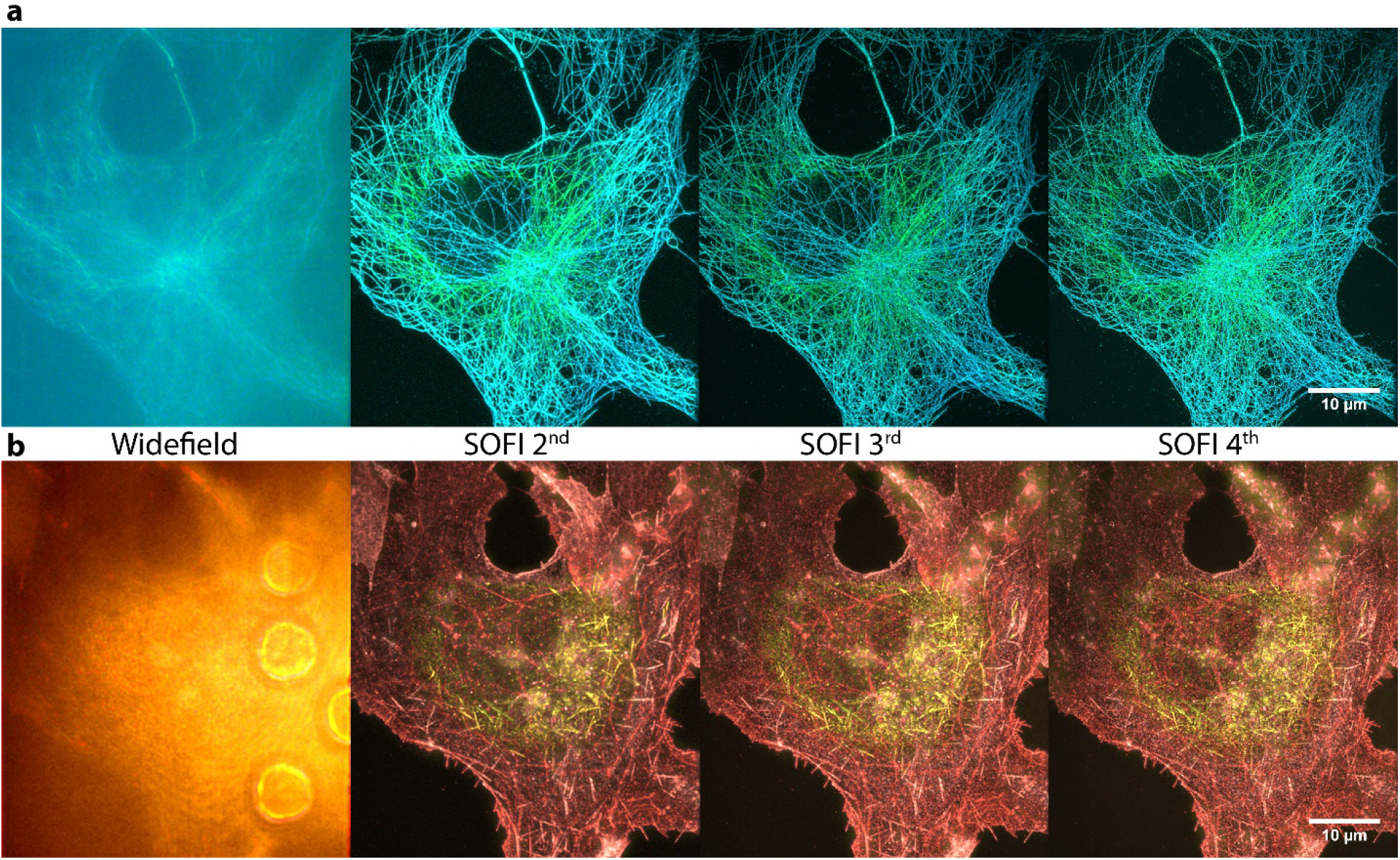
Comparison of image quality of 3D SOFI orders. SOFI orders for the two-color 3D SOFI image used in the main text Figure 3.

**Supplementary Figure 7.**
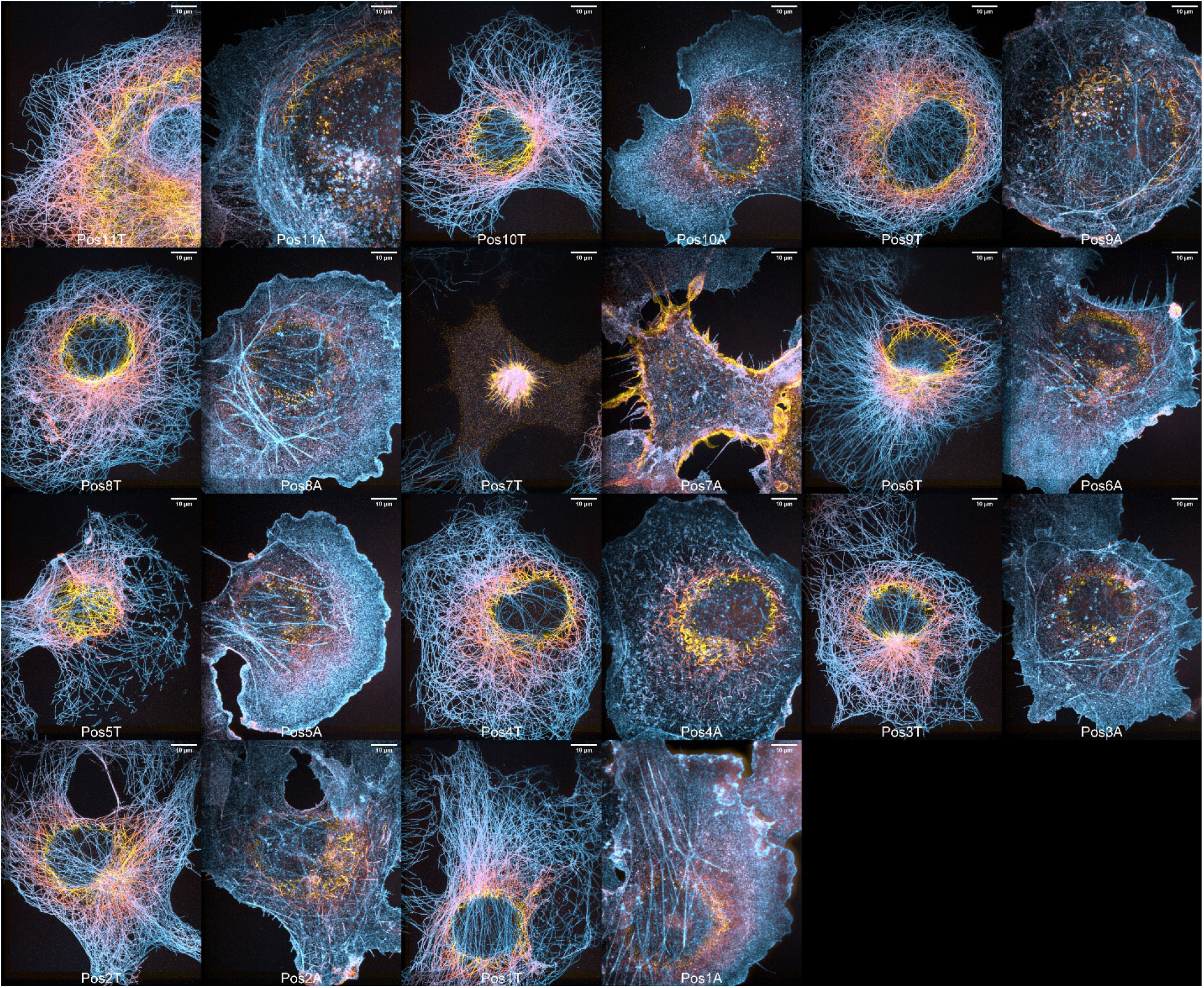
Multiple two-color 3^rd^ order 3D SOFI images. 3^rd^ order SOFI images of phalloidin-Alexa647 labeled actin (images labeled with letter A) and Abberior FLIP-565 labelled tubulin (images labeled with T). 2.45 μm axial range is represented with a color bar. Scales bars are 10 μm in all images.

**Supplementary Figure 8.**
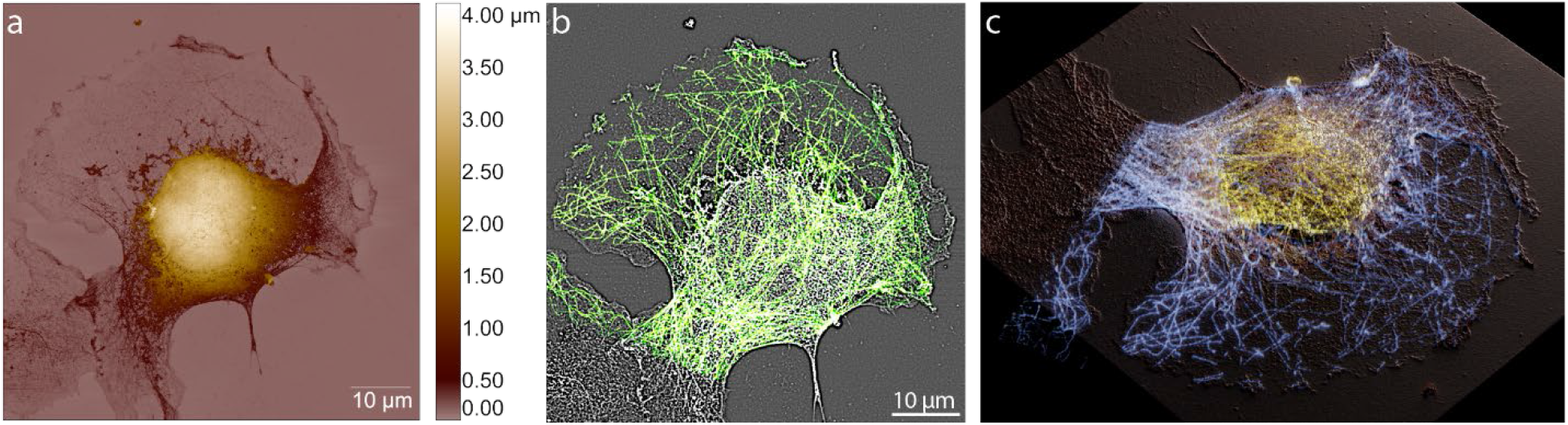
SICM topographical map of microtubules. (a) SICM image of a single cell which has part of the membrane removed by permeabilization with Triton X-100. Interestingly, this allows to reveal a preserved microtubular network of the inner part of the cell, which can be better resolved by further spatially filtering the SICM image (b) which is overlayed with a tubulin fluorescence signal from the bottom plane of 3^rd^ order 3D SOFI stack. Bandpass spatial filter with 2-8 px range was used applied on the image (a). (c) Final 3D rendering in Blender 3D software with an overlayed SOFI and SICM data.

**Supplementary Figure 9.**
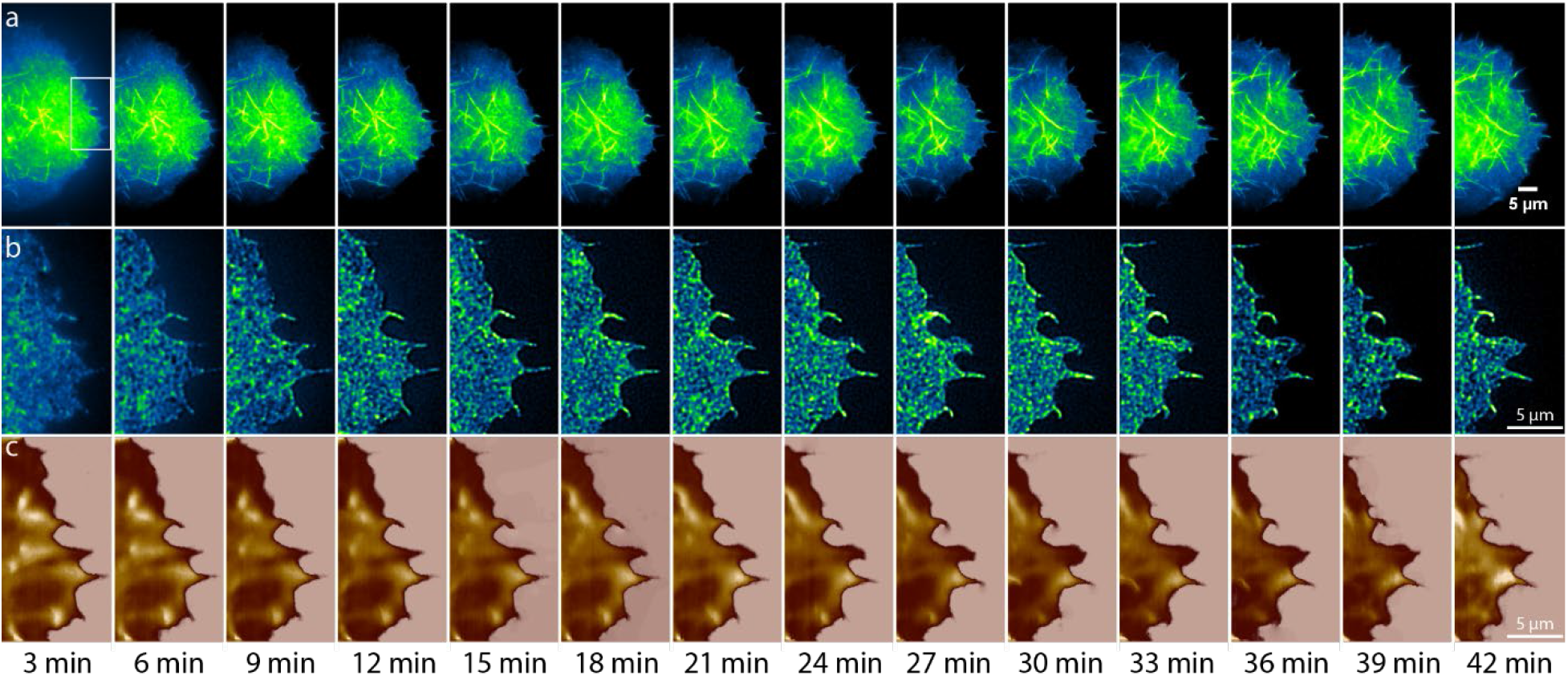
Live-cell SICM-SOFI imaging of actin of filopodia for 42 min. Acquisition was performed consequently by recording 10×20 μm SICM scans and 300 frame long fluorescence stacks for imaging photoactivated Lifeact-mEOS-2. Standard deviation image sequence (a) and 2^nd^ order SOFI (b) aligned with SICM topography images (c) are shown. COS-7 cells were transfected as described in the Methods section by using a Lifeact-mEOS-2 plasmid and imaged after 24 h in FluoroBrite medium. 200 W/cm^2^ of 561 imaging laser and 0.2 W/cm^2^ of 405 nm activation laser were used for illumination with a corresponding exposure time of 50 ms.

**Supplementary Figure 10.**
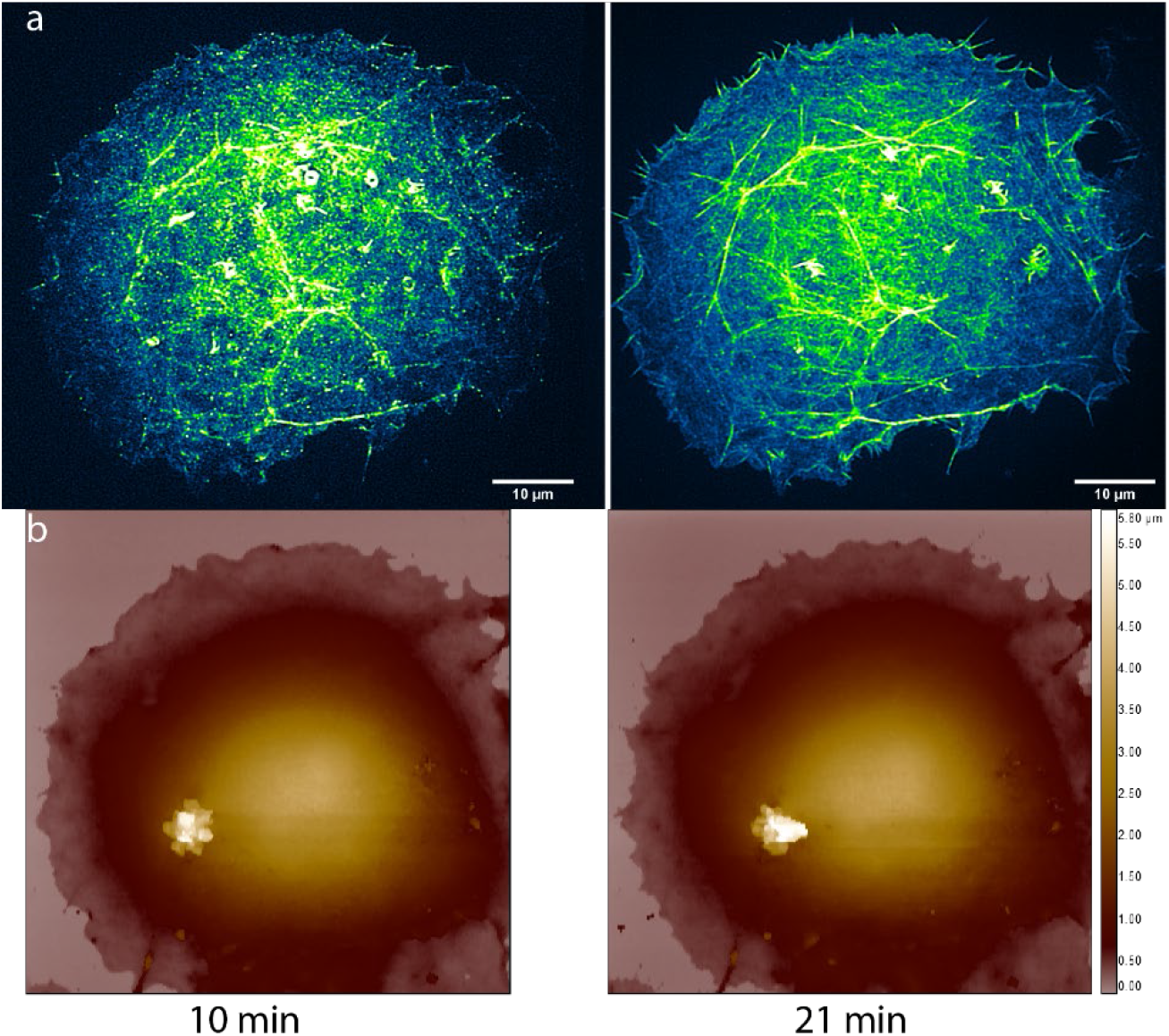
Live-cell SICM-SOFI imaging of a single cell. Acquisition was performed consequently by recording 60 x 60 μm SICM scans and 2 fluorescence stacks (300 and 1000 frames long) of photoactivated mEOS-2 for 2^nd^ order SOFI computation. 2^nd^ order SOFI (a) aligned with SICM topography images (b) are showed. COS-7 cells were transfected as described in the Methods section by using a Lifeact-mEOS-2 plasmid and imaged after 24 h in FluoroBrite medium. 200 W/cm^2^ of 561 imaging laser and 0.2 W/cm^2^ of 405 nm activation laser were used for illumination with a corresponding exposure time of 50 ms. White spot in the SICM image is most likely a particle, attached from the solution.

**Supplementary Figure 11.**
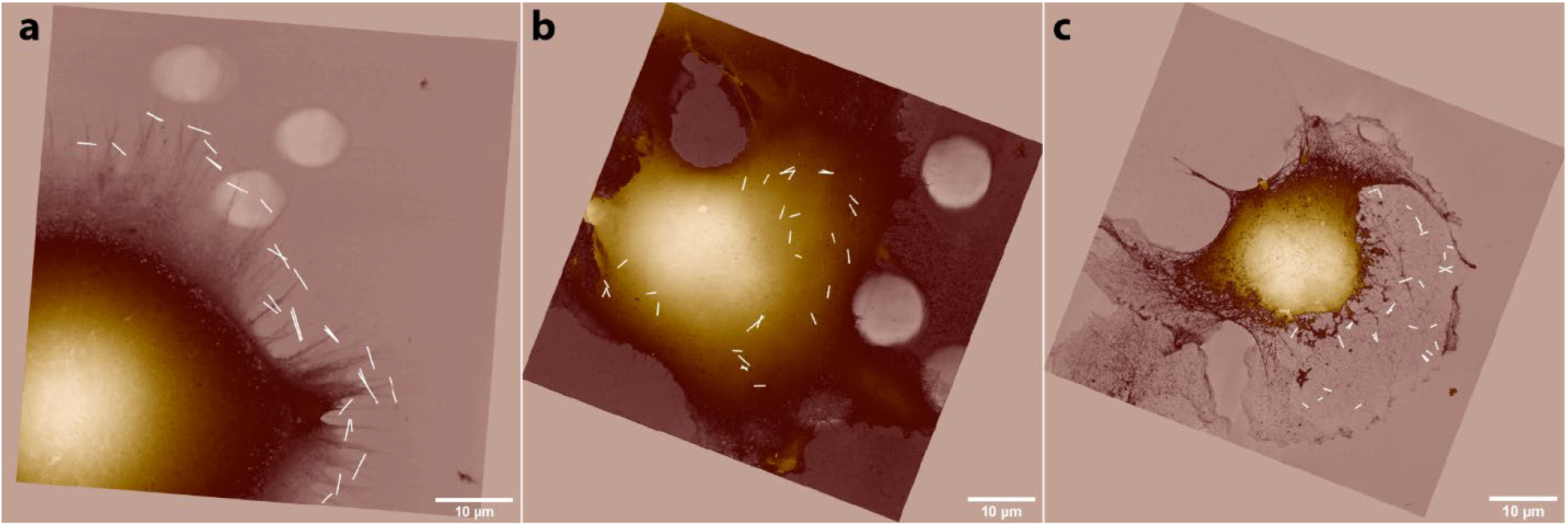
Cross sections used to calculate Pearson-correlation coefficient between different channels. For Fig 5f 1D cross sections were manually depicted by selecting the structures of interest by hand. Filopodia (a), microvilli (b) and microtubules (c) were selected.19.

**Supplementary Figure 12.**
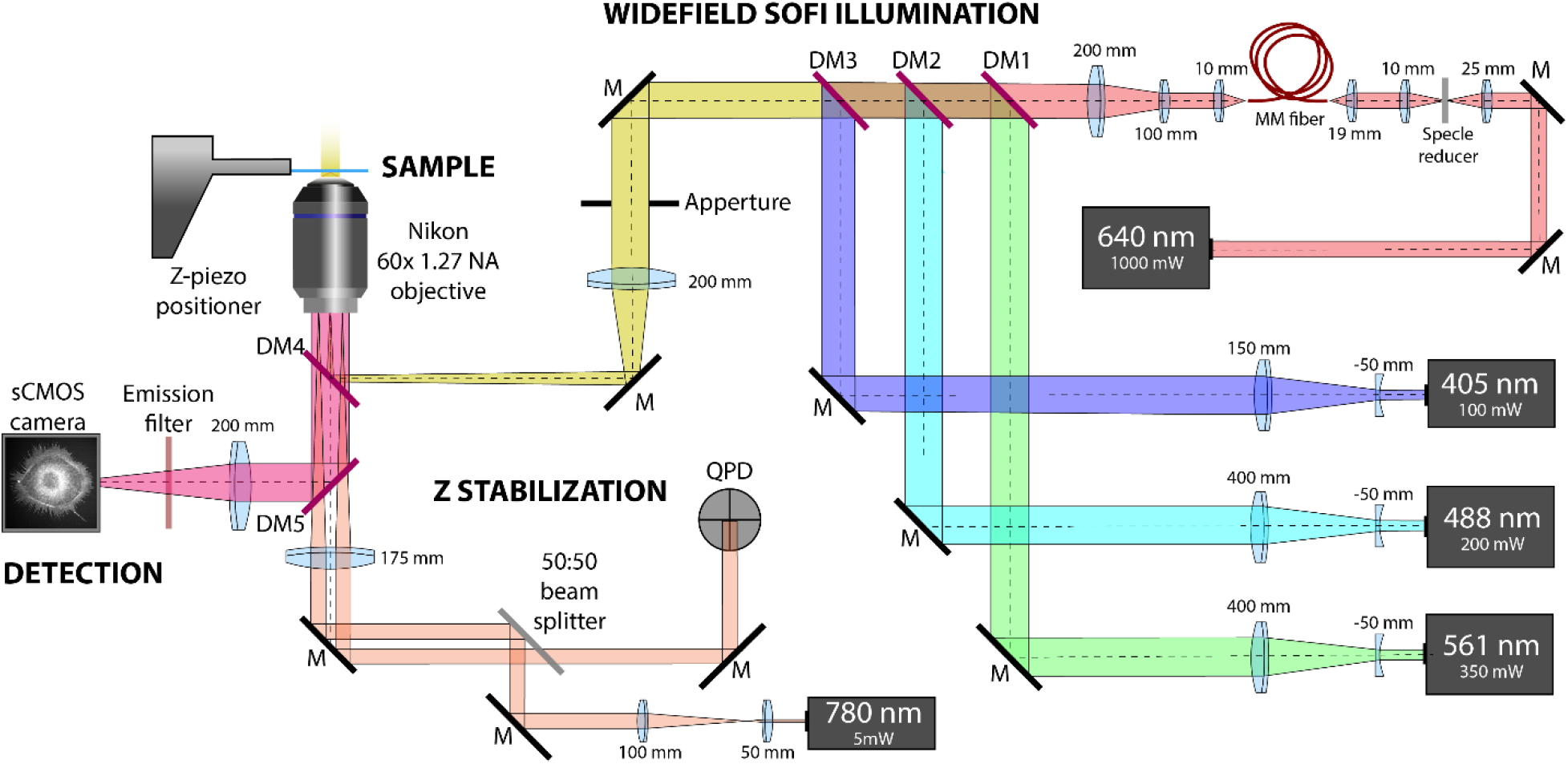
Detailed schematics of 2D SOFI setup. Schematics of the setup used for 2D SOFI imaging as described in the Methods section.

**Supplementary Figure 13.**
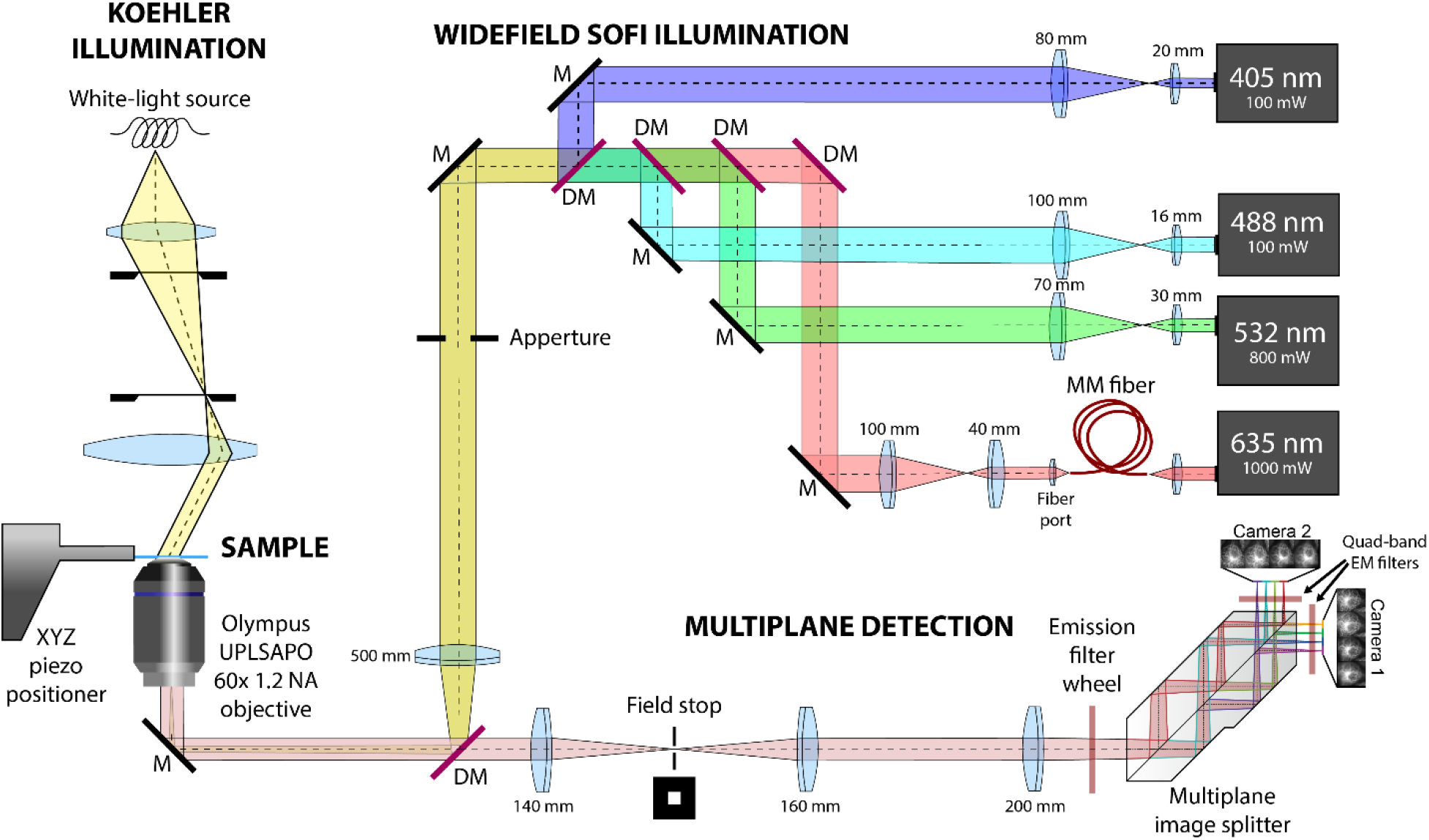
Detailed schematics of 3D SOFI setup. Schematics of the setup used for 3D SOFI imaging as described in the Methods section.

**Supplementary Figure 14.**
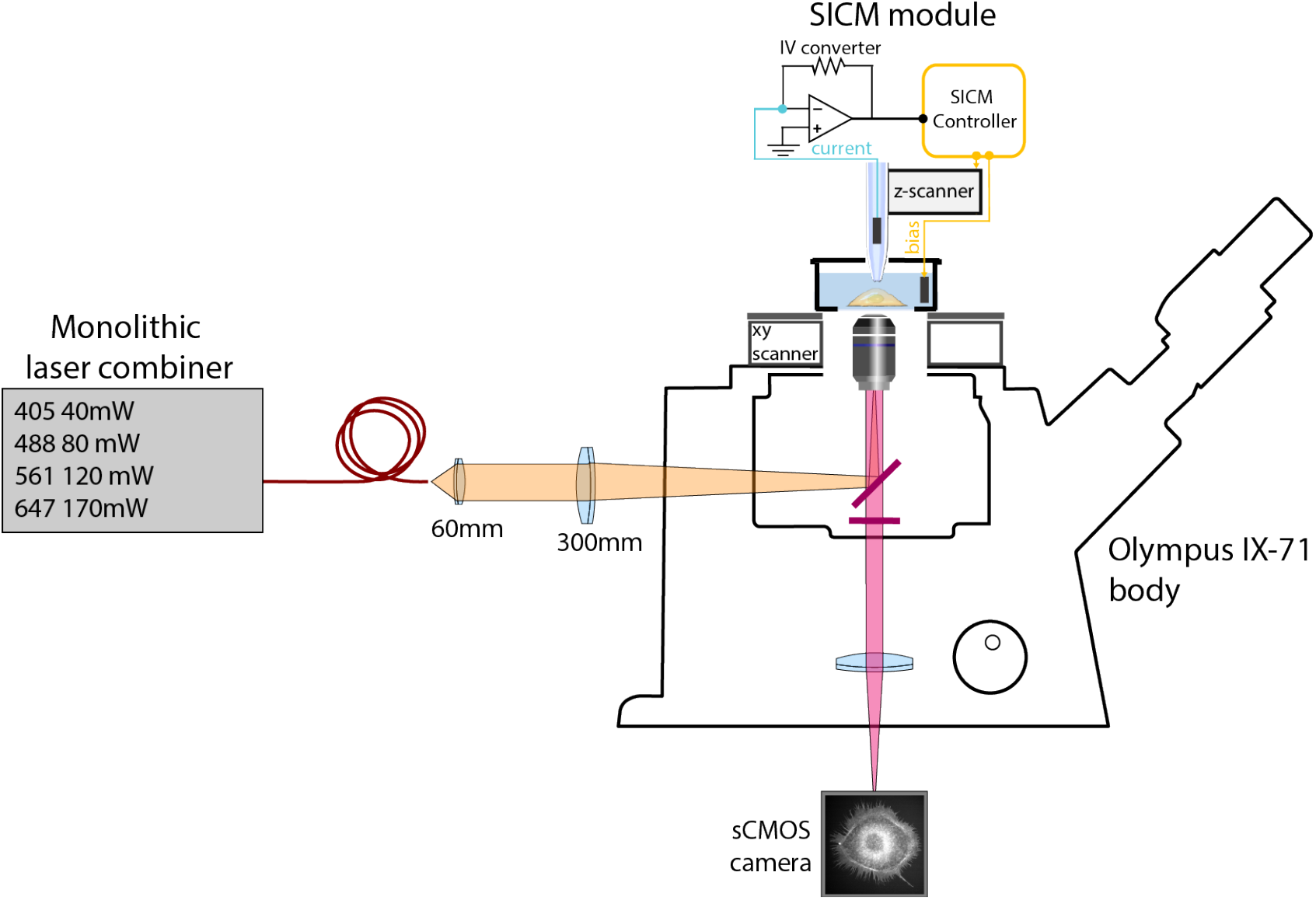
Detailed schematics of a combined SICM-SOFI setup.

**Supplementary Figure 15.**
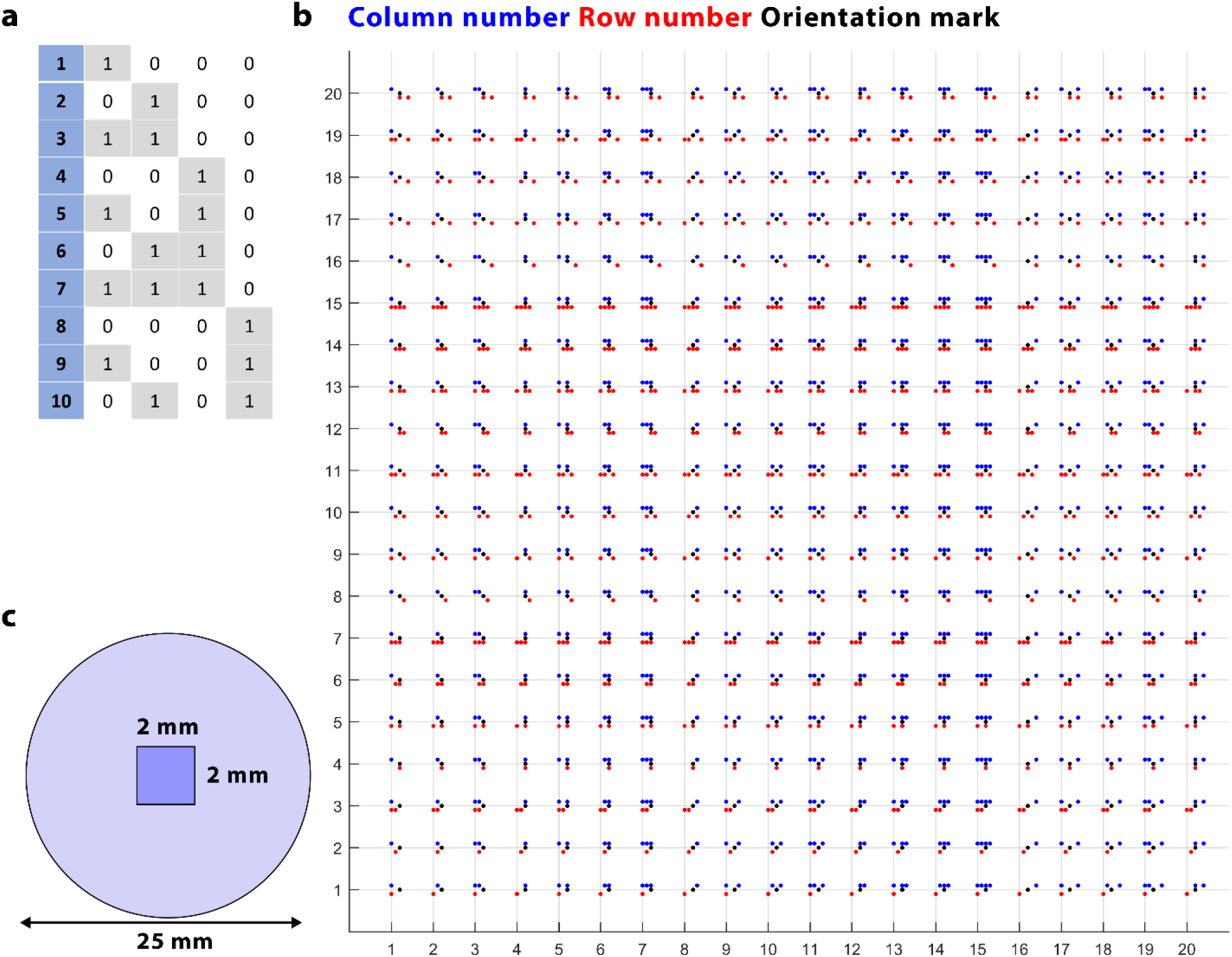
Coverslip fabrication and binary mark map generation. (a) The principle of number representation in binary values. 4 binary digits were used to represent numbers from 1 to 20. (b) The final layout of the sample map. Layout was generated with a Matlab code, by using the dec2bin function. Binary numbers were used for x and y axis. The static dot in the middle was incorporated for a better determination of the sample orientation. (c) Schematics of the dimensions of the binary digit pattern.

**Supplementary Figure 16.**
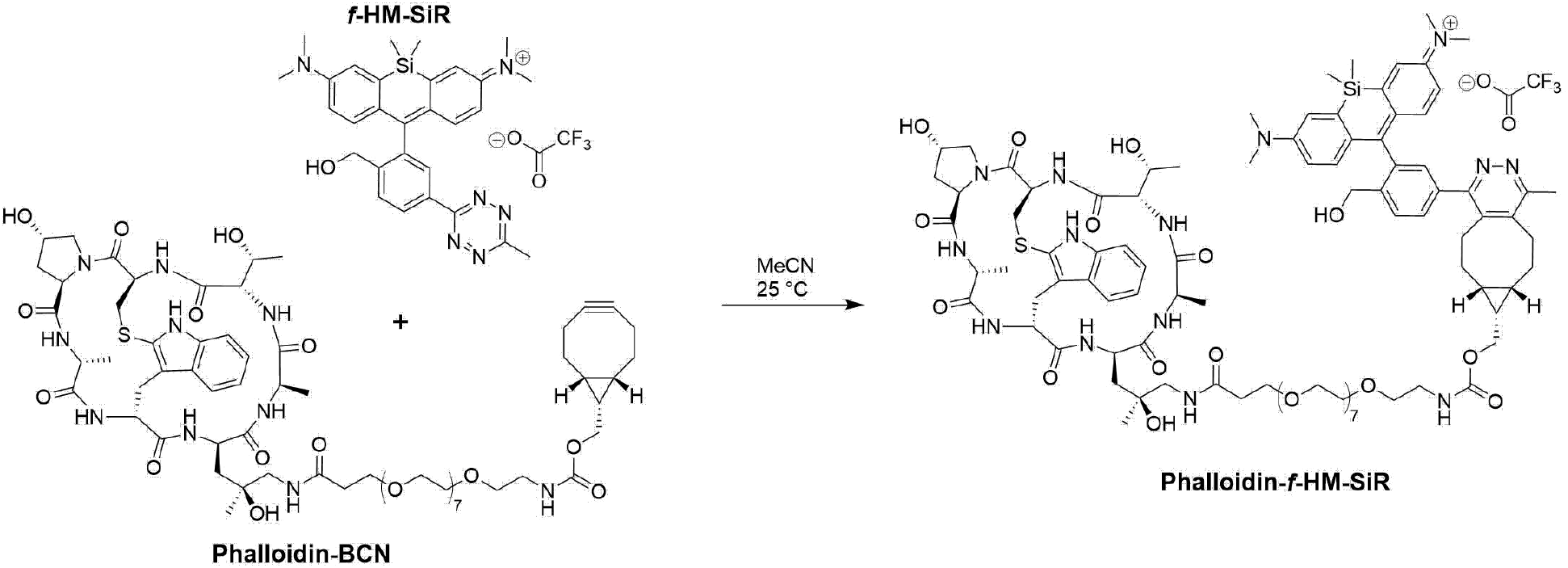
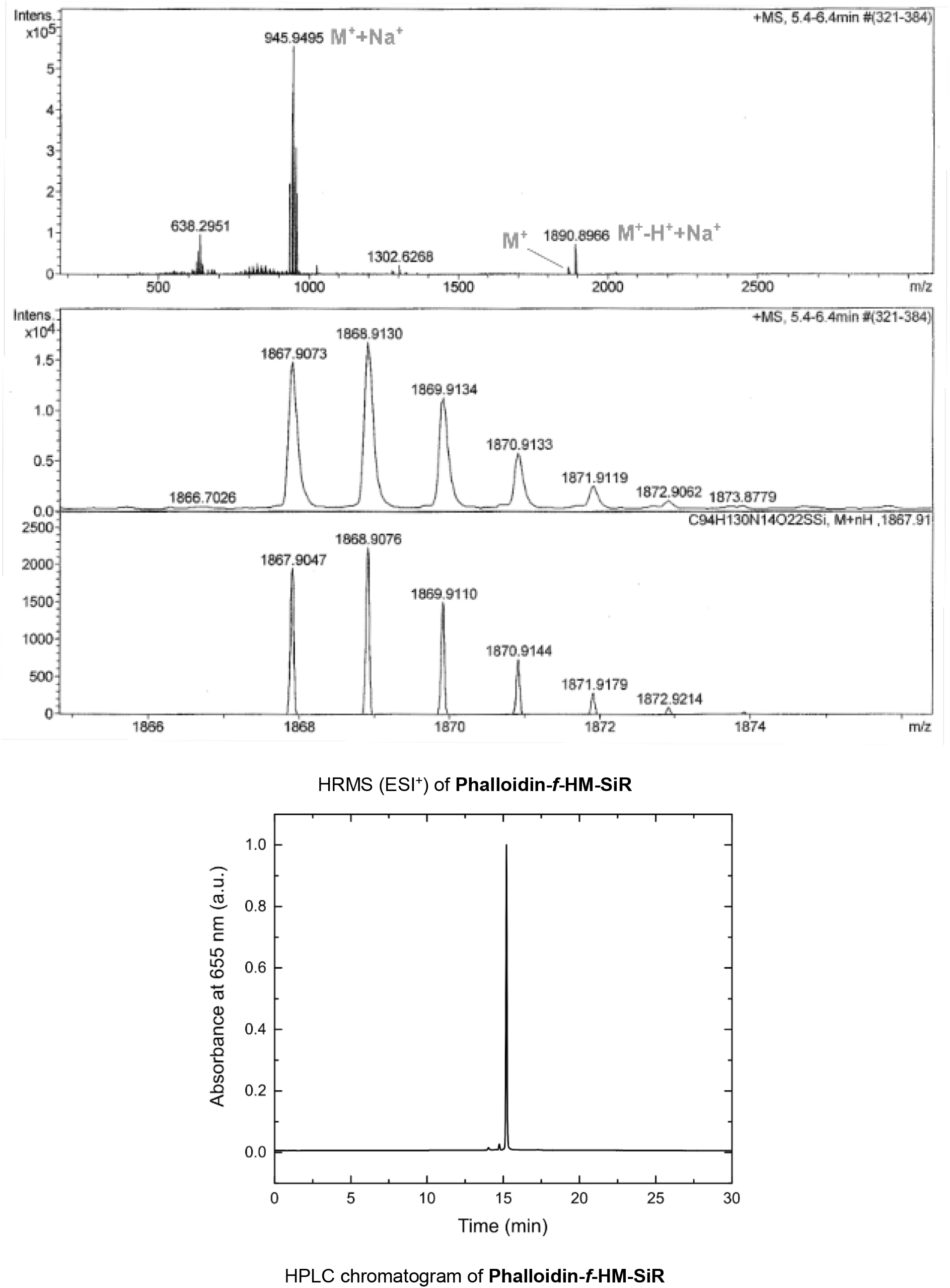
Phalloidin-f-HM-SiR chemical synthesis. Reaction scheme of Phalloidin-f-HM-SiR conjugate. **Phalloidin-*f*-HM-SiR** was synthesized from ***f*-HM-SiR**^5^ and **Phalloidin-BCN** ^6 7^, which were prepared according to literature procedures. In short: Phalloidin-BCN (46 μg) was dissolved in anhydrous MeCN (90 μL) and *f*-HM-SiR (24.8 μg, 4 μL from 10 mM stock solution) was added at room temperature. The reaction mixture was incubated for 4 h in a thermo-shaker at 700 rpm and subsequently purified by HPLC (gradient 20-90% solvent B / solvent A; in 40 min). Phalloidin-*f*-HM-SiR was afforded as blue solid and after photometric determination of the amount of substance ^8^, a 1 mM stock solution in anhydrous DMSO was prepared. HRMS (ESI^+^) m/z 1867.9047 calculated for [C_94_H_131_N_14_O_22_SSi]^+^ (M^+^), 1867.9073 found; m/z 945.4470 calculated for [C_94_H_131_N_14_NaO_22_SSi]^2+^ (M^+^+Na^+^), 945.4482 found. HPLC analytics and semi-preparative purifications were conducted on an Agilent 1100 series HPLC system. Phenomenex Luna 3 μ and 5 μ C18 reversed-phase columns were used for these purposes (Solvent A: H_2_O containing 0.1% TFA; Solvent B: MeCN containing 0.1% TFA). Collected HPLC fractions were dried by lyophilization. Mass spectrometry was performed on a Bruker microTOF-QII (for ESI-MS) mass spectrometer.

**Supplementary Figure 17.**
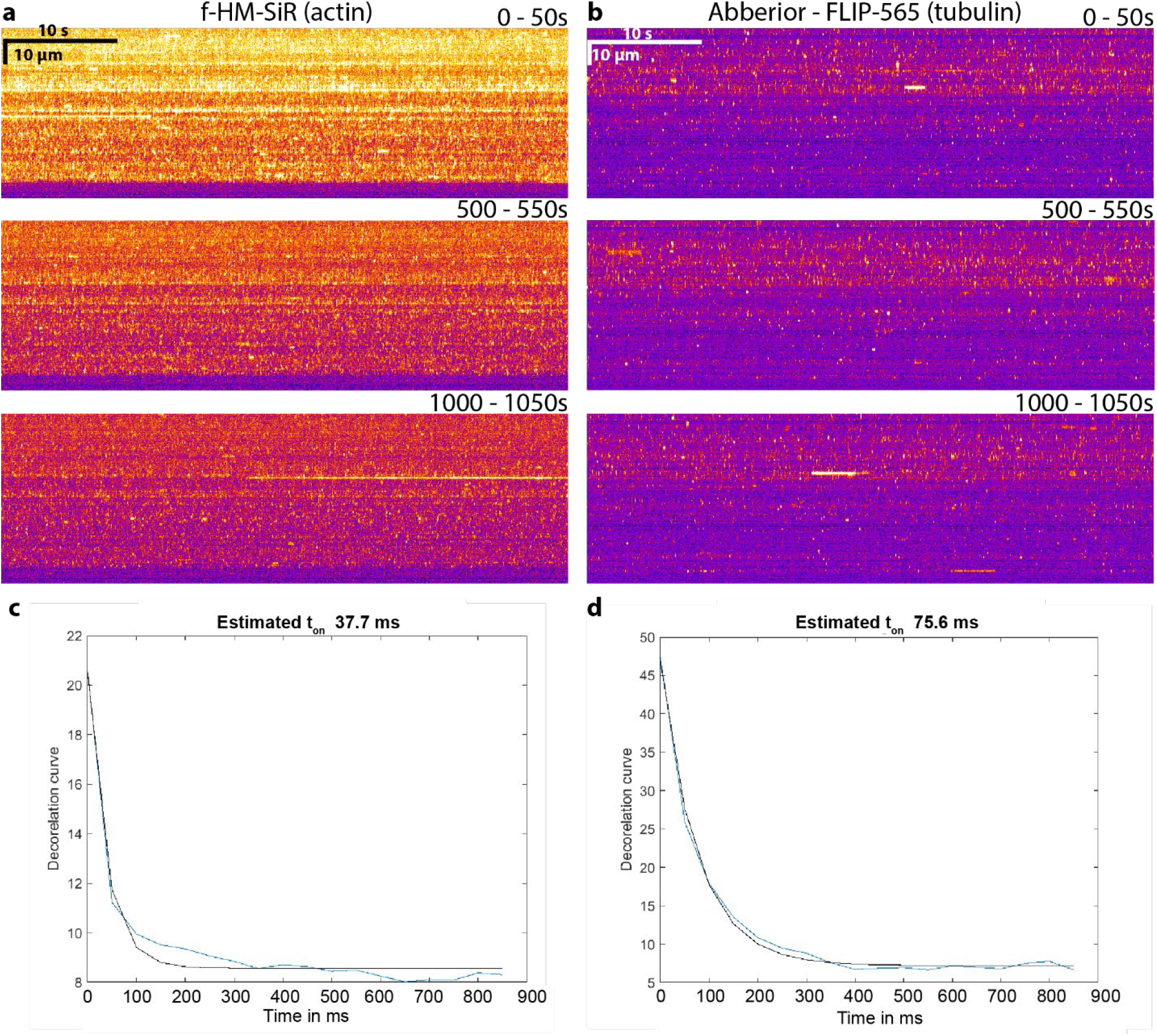
Kinetics of self-blinking dyes for high-order SOFI imaging and ON time estimation. Kymographs of signal intensity over time for f-HM-SiR (a) and Abberior-FLIP 565 (b) dyes showing the signal fluctuation over a long-term imaging. Average ON-times were also estimated by computing 2^nd^ order cumulant as a function of time lag and averaging it for subsequences of 500 frames^1^. Corresponding lag functions for stacks showed in (a-b) are showed below (c-d). Mean on time for f-HM-SiR dye was estimated to be 38.7 ms and for Abberior-FLIP 565 – 65.75 ms. However, the exposure time of 50 ms might be the limiting factor for the precision of ON-time estimation.

**Supplementary Figure 18.**
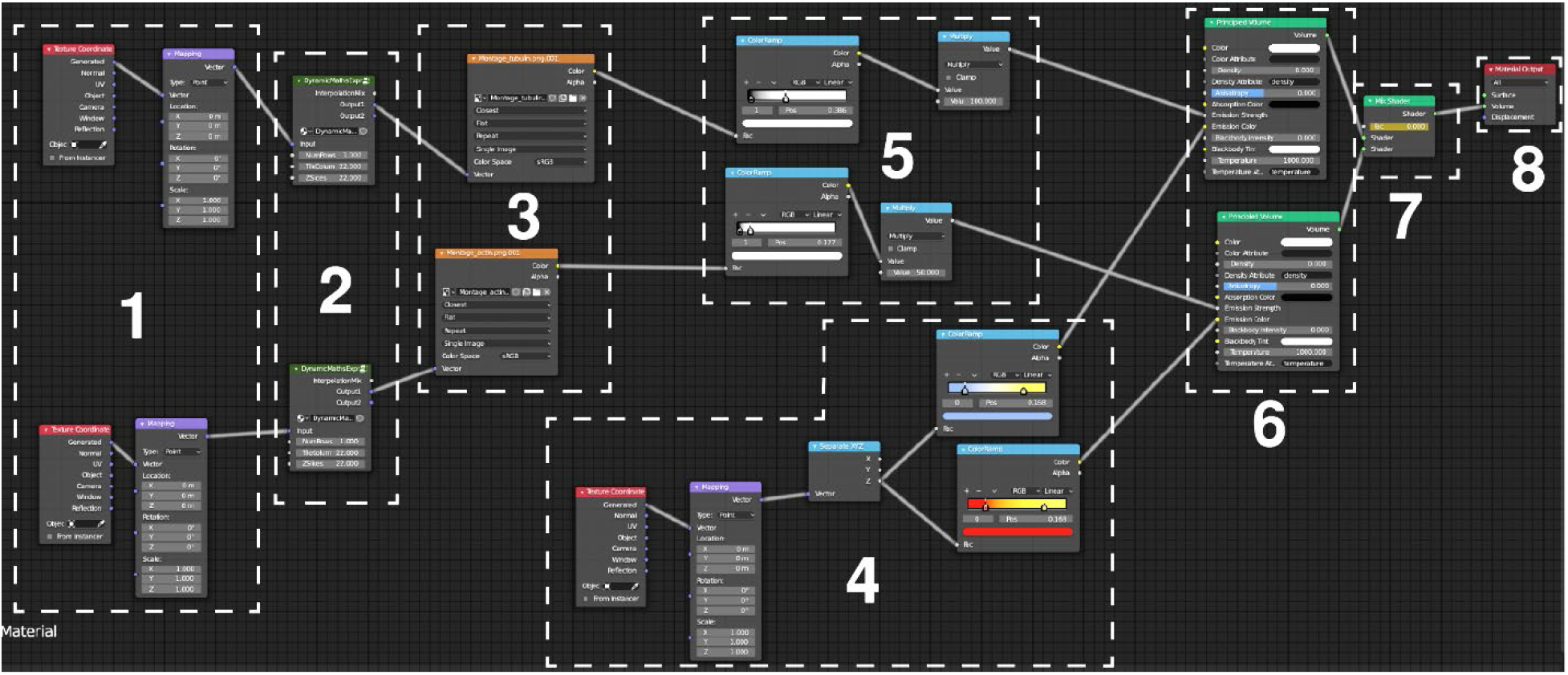
Description of final 3D SOFI data visualization in Blender 3D. A simple cube was used as a volume corresponding to the 3D SOFI volume of 2.45×60×60 μm. Two separate colors were visualized in parallel. Different parts of the shader are explained as follows: 1) Setting up the texture coordinates 2) A script node used to make a 3D volume from the stack of SOFI images. Each image is represented as voxel with dimensions determined by a 3D cube 3) Input of a stack file 4) Color ramps used to represent the height in different colors 5) Contrast and intensity adjustment 6) A principled volume shader 7) A mix shader used to switch between tubulin and actin channels 8) A volume output node.

### Lifeact-mEOS-2 sequence (ABP-tdEosFP)

Length: 5410 bp

Reference: Izeddin et al. ^9^

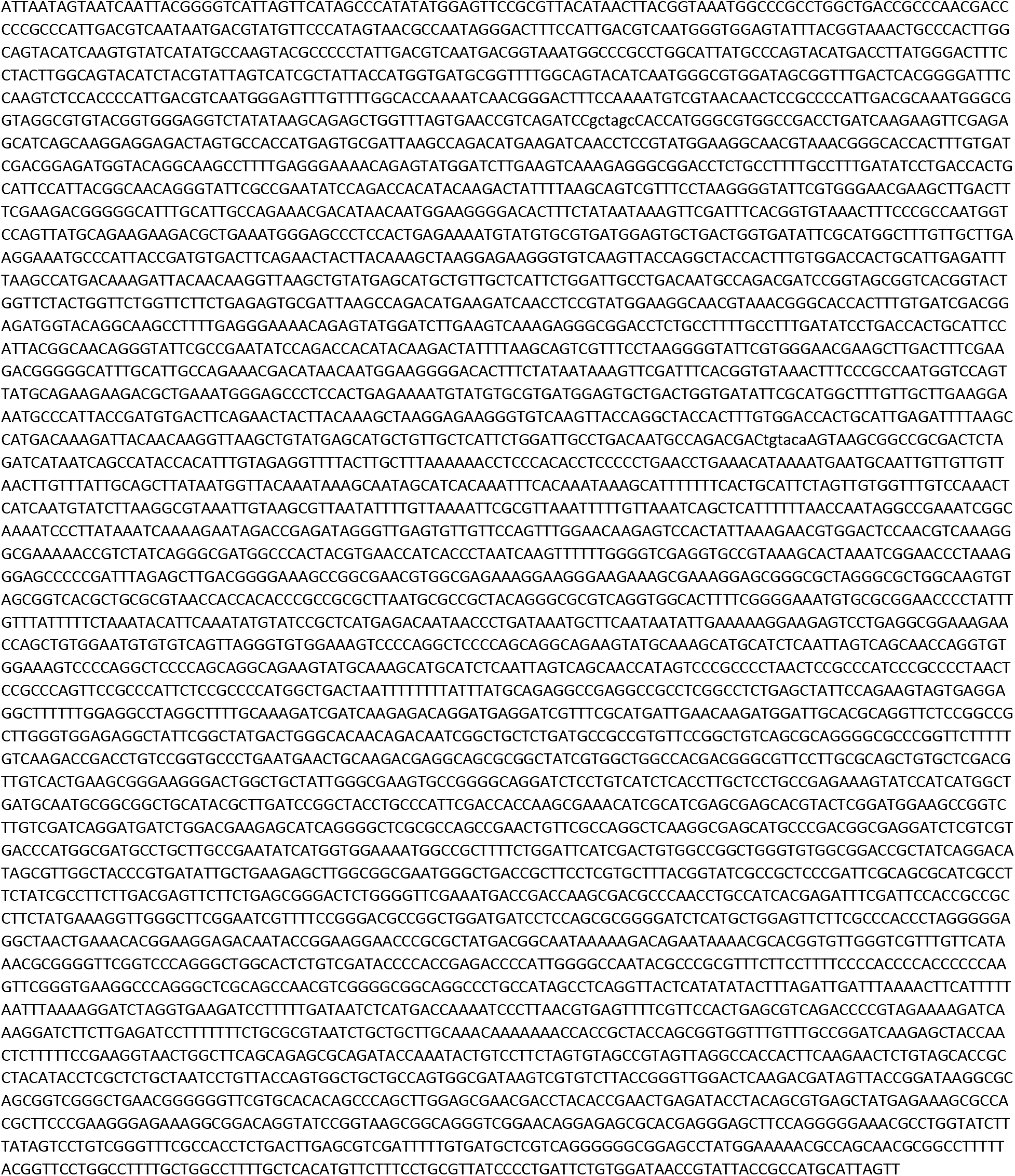

### mEos2-Alpha-Actinin-19 sequence

Length: 7384 bp

Reference: Kanchanawong et al. ^10^

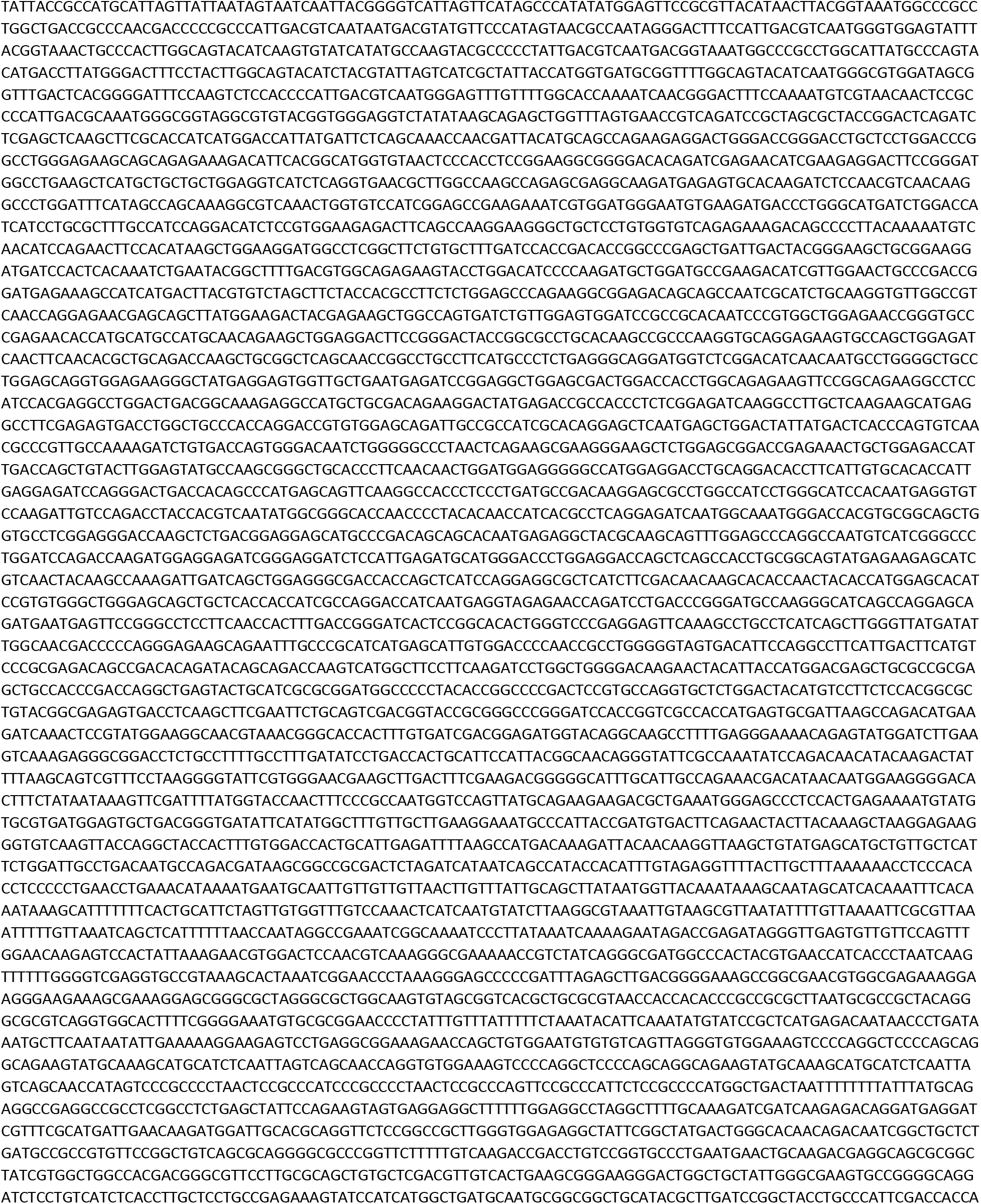
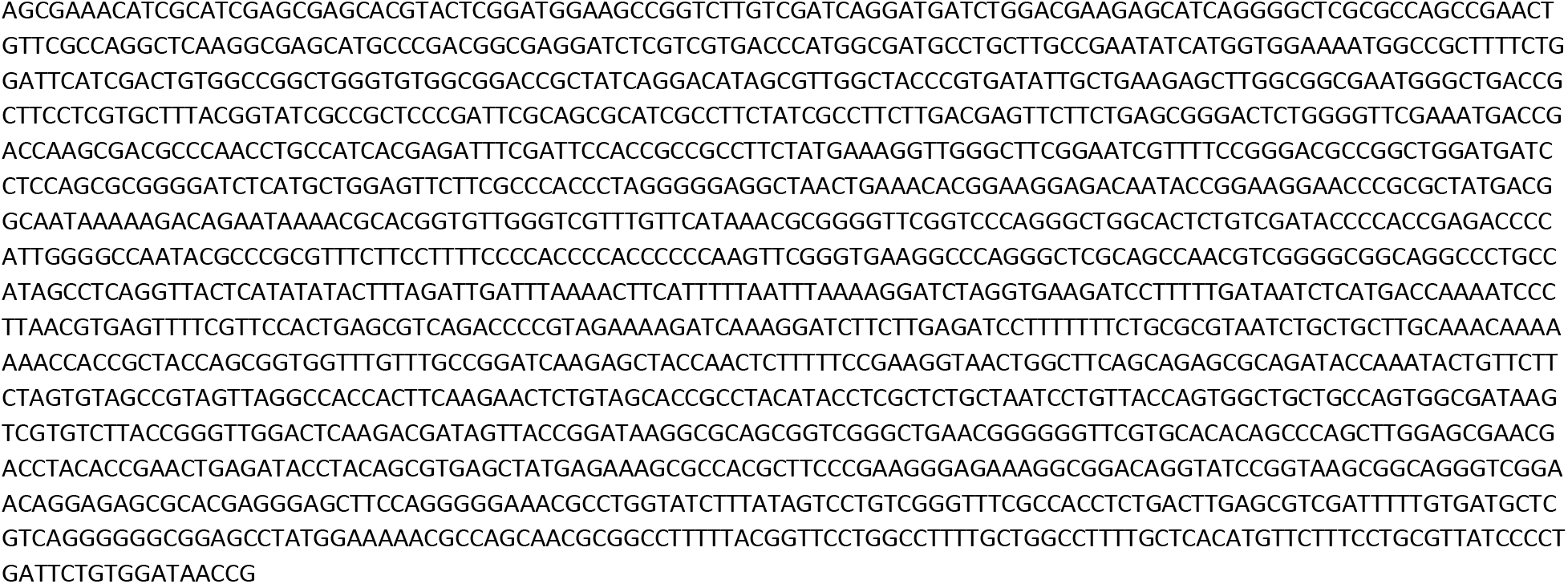

